# Evolutionary trajectory of organelle-derived nuclear DNAs in the *Triticum/Aegilops* complex species

**DOI:** 10.1101/2022.12.04.519011

**Authors:** Zhibin Zhang, Jing Zhao, Juzuo Li, Jinyang Yao, Bin Wang, Yiqiao Ma, Ning Li, Tianya Wang, Hongyan Wang, Bao Liu, Lei Gong

## Abstract

Organelle-derived nuclear DNAs, nuclear plastid DNAs (NUPTs) and nuclear mitochondrial DNAs (NUMTs), have been identified in plants. Most, if not all, genes residing in NUPTs/NUMTs (NUPGs/NUMGs) are known to be inactivated and pseudogenized. However, the role of epigenetic control in silencing NUPGs/NUMGs and the dynamic evolution of NUPTs/NUMTs with respect to organismal phylogeny remain barely explored. Based on the available nuclear and organellar genomic resources of the *Triticum/Aegilops* complex species, we investigated the evolutionary fates of NUPTs/NUMTs in terms of their epigenetic silencing and their dynamic occurrence rates in the nuclear diploid genomes and allopolyploid subgenomes. NUPTs and NUMTs possessed similar genomic atlas, including preferential integration to the transposable element-rich intergenic regions and generating sequence variations in the nuclear genome. The global transcriptional silencing of NUPGs/NUMGs with disrupted and intact open reading frames can be mainly attributed to their repressive chromatin states, namely high levels of DNA methylation and low levels of active histone modifications. Phylogenomic analyses suggested that the species-specific and gradual accumulation of NUPTs/NUMTs accompanied the speciation processes. Moreover, based on further pan-genomic analyses, we found significant subgenomic asymmetry in the NUPT/NUMT occurrence, which accumulated during allopolyploid wheat evolution. Our findings provide novel insights into the dynamic evolutionary fates of organelle-derived nuclear DNA in plants.

## Introduction

In higher plants, mitochondria and plastids originated from the endosymbiotic α-proteobacteria- and cyanobacteria-like prokaryotes, respectively **(McFadden 1999; Osteryoung and Nunnari 2003; Archibald 2015)**. Owing to the same cellular environment, extensive inter-compartmental DNA transfer among nuclear, mitochondrial, and plastid genomes occurred during the course of evolution in higher plants **(Martin, et al. 2002; Keeling and Palmer 2008; Kleine, et al. 2009; Downie and Jansen 2015)**. Among those DNA transfer events, the frequency of transfer of the chloroplast/mitochondrial DNA to the nucleus was much higher than that of the other transfer events, such as DNA transfer from the nucleus to organelle genomes or between organelle genomes **(Martin, et al. 1998; Kleine, et al. 2009; Sloan, et al. 2018; Zhao, et al. 2019)**. The nuclear plastid DNA (NUPT) and nuclear mitochondrial DNA (NUMT) refer to the organellar DNA derived from the plastids and mitochondria, respectively, which have already been incorporated in the nuclear DNA **(Leister 2005)**. NUPTs/NUMTs occur frequently and continuously; for example, more than 200 deciphered plant genomes, including *Arabidopsis*, rice, maize, and wheat, possess NUPTs/NUMTs with varying abundance **(Michalovová, et al. 2013; Zhang, et al. 2020)**. Moreover, changes in external environment and the switch of a developmental stage can lead to dramatic changes in the frequency of NUPT/NUMT occurrence in the nuclear genome **(Sheppard, et al. 2008; Caro, et al. 2010; Cheng and Ivessa 2010; Wang, Lloyd, et al. 2012)**. Several potential mechanisms underpinning the transfer and integration of plastid DNA (ptDNA) and mitochondrial DNA (mtDNA) into the nuclear genome have been posited: (*i*) direct physical association of the nucleus with the organelle, (*ii*) formation of the tubular extensions from the organelle membranes for the DNA transfer **(Leister 2005)**, and (*iii*) occurrence of double-strand breaks (DSBs) in the nuclear genome, facilitating ptDNA /mtDNA integration between those breaks by the non-homologous end joining pathway or through homologous recombination **(Kleine, et al. 2009; Hazkani-Covo, et al. 2010; Portugez, et al. 2018)**.

Several studies have reported that after the integration into the nuclear genome, NUPTs/NUMTs could generate nucleotide mutations **(Huang, et al. 2005)**, amplified together with the hosting transposable elements (TEs) **(VanBuren and Ming 2013)**, or get fragmented because of TE insertion **(Michalovová, et al. 2013)**. The foregoing processes are prone to inducing interruptions in the open reading frames (ORFs) of those organellar genes residing in NUPTs/NUMTs (NUPGs/NUMGs). However, the inactivation and pseudogenization of NUPGs/NUMGs after their integration into the nuclear genome is still an underexplored area of study **(Park, et al. 2020)**. Considering the importance of epigenetic modifications in gene activity **(Feng and Jacobsen 2011; Zhang, et al. 2018)**, the role of epigenetic silencing in the transcriptional inactivation of NUPGs/NUMGs should be explored. Three exemplary studies on the inactivation of NUPGs/NUMGs independently reported NUPTs/NUMTs as the alien nuclear genetic materials (like TEs) to be silenced through genomic defense **(Zhang, et al. 2020)**, the important role of DNA methylation against NUPTs to maintain genome stability **(Yoshida, et al. 2019)**, and the possible role of epigenetic modification in the silencing of a NUMT fragment in *Arabidopsis* **(Fields, et al. 2022)**. However, these studies did not compare the DNA methylation patterns of NUPGs/NUMGs with the indigenous nuclear genes; moreover, the generality of their conclusions remained unclear.

Besides diploid divergence and speciation, the ubiquitous whole-genome duplication or polyploidization has played a pivotal role in the evolution and speciation of angiosperms **(Adams and Wendel 2005; Jiao, et al. 2011; Van de Peer, et al. 2017)**. Theoretically, in a polyploid plant, multiple nuclear subgenomes are merged into the same nucleus; thus, more sites are available for NUPTs/NUMTs.Based on the established subgenome dominance embodied as asymmetric expression, epigenetic modification, structural variation and TE dynamics **(Pont and Salse 2017; Bird, et al. 2018; Li, et al. 2021)**, whether the evolutionary dynamics of NUPTs/NUMTs is related to these features of subgenome asymmetry poses an intriguing question.

The *Triticum/Aegilops* complex consists of 31 species including 14 diploid, 11 allotetraploid, and 6 allohexaploid species **(Ogihara, et al. 2016)**. Around 7 million years ago (MYA), ancient *Triticum* and *Aegilops* species diverged into two diploid lineages, namely A- and B-lineages; thereafter, the D-lineage species were derived from the homoploid hybridization between A- and B-lineage species. Common wheat (*Triticum aestivum*) harboring three distinct subgenomes, A (from *T. urartu* in A lineage), B (from an unknown species related to *Aegilops speltoides* in B lineage), and D (from *Ae. tauschii* in D lineage), was eventually developed via two distinct rounds of allopolyploidization events **(Marcussen, et al. 2014; Levy and Feldman 2022; Li, et al. 2022; Xiao, et al. 2022)**. Similarly, the *Triticum/Aegilops* complex species encompass a reticulate evolutionary trajectory involving diploid speciation, allopolyploidization, and crop domestication and improvement. Moreover, since a series of high-quality nuclear and organelle genome assemblies in the *Triticum/Aegilops* complex species have been recently published **(Avni, et al. 2017; Luo, et al. 2017; Consortium, et al. 2018; Ling, et al. 2018; Maccaferri, et al. 2019; Walkowiak, et al. 2020; Wang, et al. 2020; Fu 2021; Li, et al. 2022)**, these genomic resources established the basis for systematic investigation of the dynamic evolution of NUPTs/NUMTs at the phylogenomic scale **(Liang, et al. 2018)**.

In this study, we investigated the evolutionary fates of NUPTs/NUMTs by delineating the genome-wide atlas of NUPTs/NUMTs in the diploid and allopolyploid *Triticum/Aegilops* species. Using common wheat as a reference, we also characterized the mutational features, expression profiles, and the role of epigenetic modification in the silencing of NUPGs/NUMGs. We constructed a phylogenomic- and pan-genomic-based pipeline to analyze the evolution pattern of the genomic/subgenomic NUPTs/NUMTs during the diploid speciation and allopolyploid evolution of the *Triticum/Aegilops* species. Our results provide novel insights into the dynamic evolutionary fates of organelle-derived nuclear DNAs in plants.

## Results

### The landscapes of NUPTs/NUMTs in the *Triticum*/*Aegilops* complex species

The genomic/subgenomic sequences of interest in the *Triticum*/*Aegilops* complex species were identified. A total of 1,860–2,954 NUPT (1.26–3.35 Mb; 0.026%– 0.057%) and 2,440–4,787 NUMT (3.56–8.12 Mb; 0.084%–0.180%) high-confidence sequences were identified (**Figure 1B and C and Figure S3B and C**; **see Materials and Methods**), and NUMT proportion was higher than that of NUPTs (**Figure 1D**; Mann–Whitney U test, *p* value < 0.001). The proportion of NUPTs and NUMTs varied across different species lineages: NUPTs, D lineage (2,585–2,954, 2.00–3.35 Mb) > B lineage (2,018–2,213, 1.33–1.95 Mb) ≈ A lineage (1,860–2,226, 1.27–1.78 Mb); NUMTs, D lineage (3,250–4,787, 5.39–8.21 Mb) ≈ B lineage (except *Ae. speltoides*; 3,078–3,142, 6.71–6.98 Mb) > A lineage (2,367–2,818, 4.29–4.81 Mb) (**Figure 1B and C and Figure S3B and C**).

**Figure 1.**
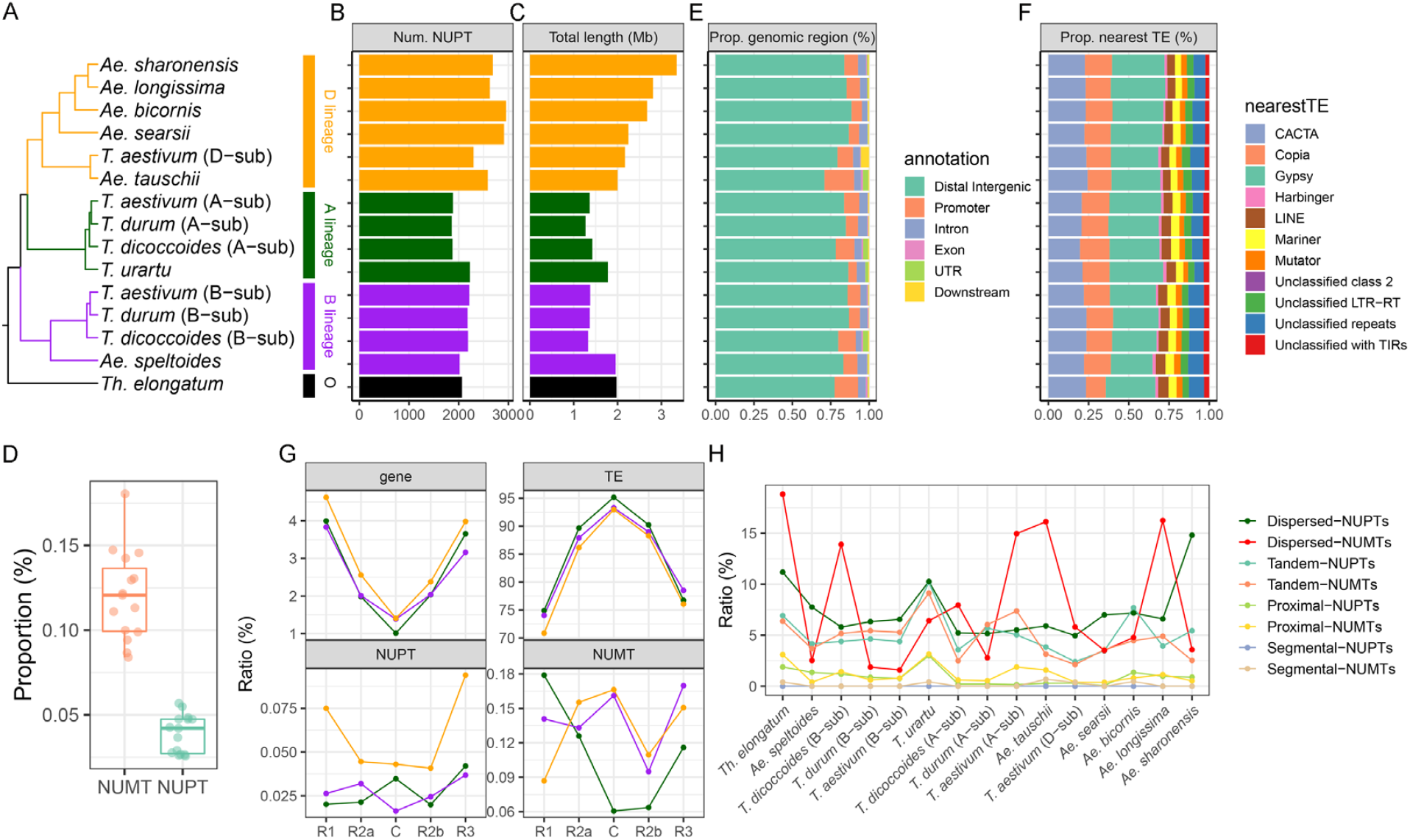
Genomic landscape of NUPTs/NUMTs in the *Triticum*/*Aegilops* complex species. **(A)** Phylogenetic tree topology in the *Triticum*/*Aegilops* complex species from the study of Li *et al*. Whole-genome statistics of NUPTs including **(B)** numbers, **(C)** total length, **(E)** the proportion of the distribution of different genomic features, and **(F)** the proportion of nearest transposon types. **(D)** The proportion of the total length of NUPTs and NUMTs; each point represents a given genome (diploid) or subgenome (allopolyploid). **(G)** The relative proportion of genes, transposable elements, and NUPTs/NUMTs in five different chromosome regions of the IWGSC RefSeq 1.0 genome; the division of chromosome regions is based on the work of Consortium *et al*. **(H)** The proportion of different types of duplicated NUPTs/NUMTs in each genome/subgenome, including dispersed, tandem, proximal, and segmental NUPTs/NUMTs. The corresponding genomic characteristics of NUMTs are shown in **Figure S3**.

The genomic distribution of both NUPTs and NUMTs was conserved across the genomes/subgenomes. The majority of NUPTs/NUMTs were located in the intergenic regions (70.9%–88.6% for NUPTs; 76.3%–90.0% for NUMTs) and were especially enriched near the *Gypsy* (25.8%–33.8% for NUPTs; 29.3%–36.0% for NUMTs), *CACTA* (19.7%–24.4% for NUPTs; 18.9%–22.6% for NUMTs), and *Copia* (12.3%– 18.4% for NUPTs; 12.5%–19.0% for NUMTs) TEs (**Figure 1E–F**). The distribution of NUPTs/NUMTs, relative to the protein-coding genes and TEs present in the chromosomes, was further compared (**Figure 1G**) based on the IWGSC RefSeq 1.0 genome (*T. aestivum* Chinese Spring variety). The gene density gradually decreased from the telomeric region to the centromeric region, whereas TEs exhibited opposite trends in all three subgenomes (**Figure 1G**). Intriguingly, neither NUPTs nor NUMTs showed a distribution similar to that of genes or TEs. The NUPT/NUMT distribution patterns within different subgenomes were distinct, suggesting that large-scale subgenomic/species-specific integration of the plastid/mitochondrial DNA happened during the evolution of the *Triticum*/*Aegilops* complex species.

Additionally, the occurrence and the extent of the second amplification/duplication events of NUPTs/NUMTs were investigated. Intriguingly, similar to other indigenous genic duplication events, the results showed a large scale of endo-nuclear replication events for both NUPTs and NUMTs (**see Materials and Methods**). A total of 169–544 (9.0%–23.4%) NUPTs and 165–920 (6.6%–28.7%) NUMTs had at least one duplication event, where the major duplication classes were of dispersed duplication (43.8%–70.0% for NUPTs; 20.7%–72.9% for NUMTs) and tandem duplication (25.7%–51.5% for NUPTs; 14.6%–68.9% for NUMTs), followed by proximal duplication (1.5%–12.8% for NUPTs; 4.6%–16.5% for NUMTs).

Segmental duplication events occurred only for five species/subgenomes, 4.4% for NUMTs and none for NUPTs. The endo-nuclear replication results of NUPTs/NUMTs suggested their potential genetic effects on reshaping the nuclear genomic structure in the *Triticum*/*Aegilops* complex species.

### Genetic variations resulting in the loss of the coding ability of NUPGs/NUMGs

After aligning to the corresponding organelle genomes, all NUPTs in each of the genome/subgenome covered the whole chloroplast genome regions, from 2x depth of inverted repeat region b (IRb) in *Triticum dicoccoides* B-subgenome to 65x depth of large single-copy (LSC) region in *Aegilops sharonensis* for genetic variability (**Figure 2A**). The results suggested that the ubiquitous chloroplast DNA (cpDNA) sequences could transfer into the nuclear genome. During genomic evolution, NUPTs/NUMTs could generate a wide range of genetic variations, such as single-nucleotide polymorphisms (SNPs) and insertion/deletions (InDels). A detailed analysis of respective SNPs and InDels in NUPTs among all the genomes/subgenomes with reference to their indigenous chloroplast genomic sequences revealed the following: (*i*) a total of 12.7%–21.5% SNP sites (70% non-redundant SNPs in total), among which respective density ranged from 32 SNPs/kb in IRb in *Thinopyrum elongatum* to 314 SNPs/kb in LSC regions in *Ae. bicornis* (**Figure 2B**); (*ii*) a total of 1.6%–15.4% InDels (35.0% non-redundant InDels in total), among which the respective density ranged from 2 InDels/kb of IRa (inverted repeat region a) regions in *T. durum* B-subgenome to 274 InDels/kb in LSC regions in *Ae. sharonensis* (**Figure 2C**); (*iii*) SNPs and InDels were highly similar among NUPTs of all the genomes/subgenomes in terms of their types, transitions, and transversions for SNPs and InDels of different lengths. For SNPs, the proportion of transitions (especially G to A and C to T) was larger than that of transversions, which is consistent with the results of a previous study **(Noutsos, et al. 2005)**. For InDels, the proportions of 1-bp variations (both insertion and deletion) were significantly overwhelming compared with the other length classes (**Figure 2D–E**); (*iv*) In NUPTs of all the genomes/subgenomes, the proportions of conserved SNPs (0.10%) and InDels (0.05%) were very low, and 29.1% of SNPs and 55.1% InDels were genome-specific (**Figure 2F and G**); (*v*) mutations within the same genome/subgenome origination (A, B, D, and S) were highly shared, resulting in the phylogenomic-mimic clustering pattern, especially for SNPs (**Figure 2F and I**). NUMTs exhibited similar mutation patterns as NUPTs; moreover, they had certain conserved genomic variations as well as a high proportion of species-specific variations (**Figure S4**).

**Figure 2.**
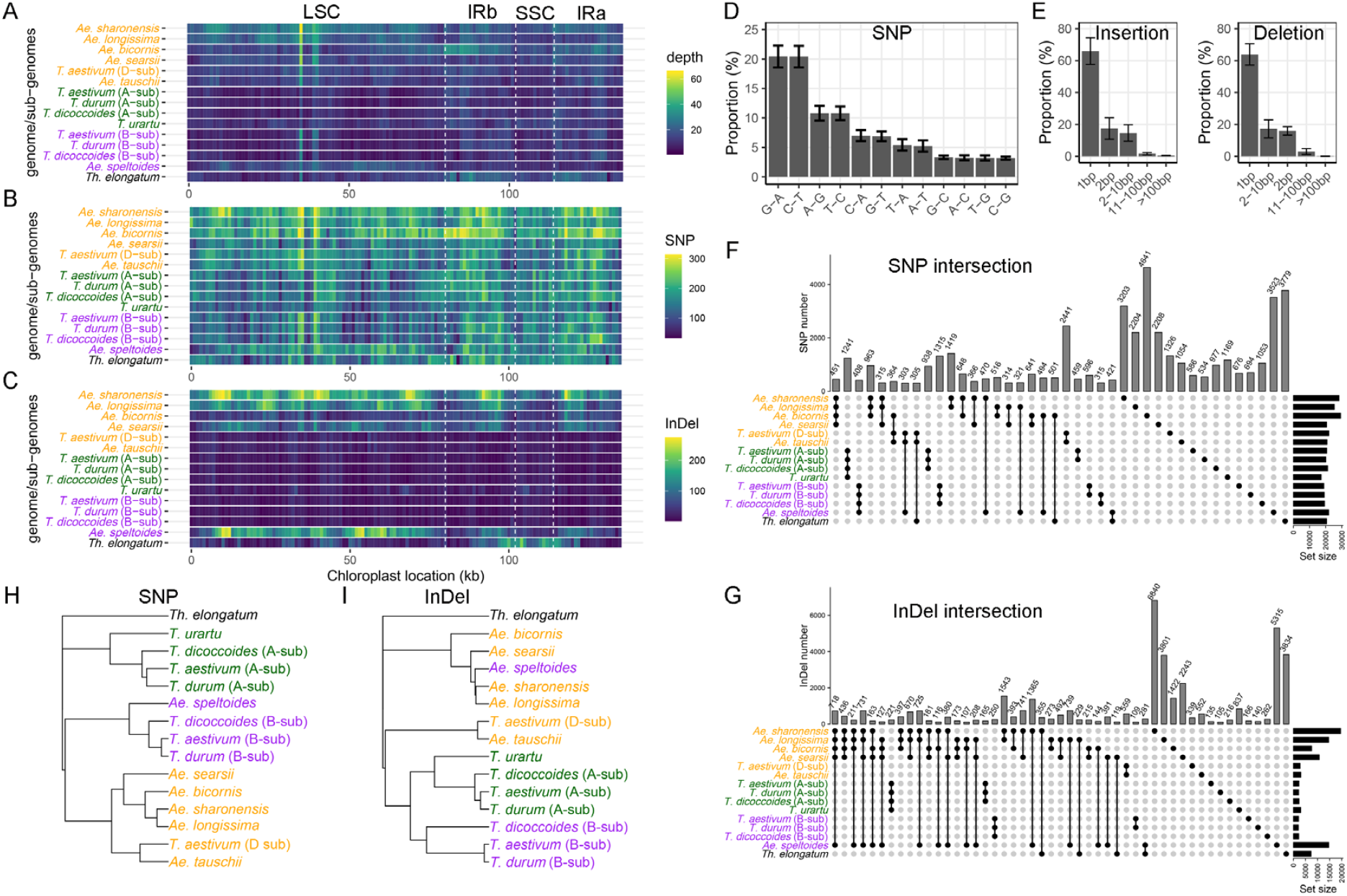
Characteristics of genetic variations in NUPTs compared with the chloroplast genome. The density feature of NUPTs in different regions of the chloroplast genome based on non-overlapping 1-kb windows, including **(A)** insertion frequency (depth), **(B)** single nuclear polymorphism (SNP) density, and **(C)** InDel density. NUPTs were first aligned to the chloroplast genomes and then examined for each feature. LSC: large single-copy region; SSC: small single-copy region; IRa: repeat region a; and IRb: repeat region b. The vertical dashed lines represent the junction of the above regions. **(D)** The proportion of different types of SNPs. The error bars indicate the standard deviation among different genomes/subgenomes. **(E)** The proportion of different types of insertions and deletions. **(F)–(G)** The UpSet plot based on the intersection matrix of SNPs **(F)** and InDels **(G)** in each variation site among genomes/subgenomes. **(H)–(I)** The neighbor-joining tree topology based on the intersection matrix of SNPs **(H)** and InDels **(I)** used in **(F)** and **(G)**. The corresponding information on NUMTs is shown in **Figure S4**.

Based on the SNP/InDel results, the genetic fate of the NUPG/NUMG ORF disruptions, via fragmentation and premature and frameshift mutations, was explored to determine the intact and disrupted ORFs based on the maintenance or loss of the original coding ability of ORFs, respectively. Genes with intact and disrupted ORFs were named as NUPGs/NUMGs and d-NUPGs/NUMGs, respectively. All organellar genes were analyzed for their susceptibility to integration in the nuclear genome. The NUPT/NUMT alignment with respective chloroplast/mitochondrial genomes identified 234 (*T. dicoccoides* B-subgenome)–1,170 (*Ae. sharonensis*) and 152 (*Ae. speltoides*)–395 (*Ae. bicornis*) sequences harboring both NUPGs/NUMGs and d-NUPGs/NUMGs. Among them, d-NUPGs ranged from 216 (45.3%; *T. urartu*) to 890 (76.1%; *Ae. sharonensis*), and d-NUMGs ranged from 173 (48.5%; *Ae. tauschii*) to 358 (90.6%; *Ae. bicornis*) (**Figure 3A and Figure S5A**). Among the proportions occupied by d-NUPGs, *Sitopsis* genomes, except that of *Ae searsii*, were the largest (69.1%–76.0%), followed by A- (except *T. urartu*, 56.7%–60.0%), B-(47.8%– 54.7%), and D (47.5%–50.8%)-genomes/subgenomes (**Figure 3A)**. The proportions of d-NUMGs in the different genomes/subgenomes existed in a similar order as that of NUPGs but with an overall higher probability of disrupted ORFs (except *Ae. searsii* and *T. dicoccoides* A subgenomes) (**Figure S5A**). The frequency of the intact NUPG/NUMG ORFs was determined for different organellar genes (**Figure 3B and Figure S5B**). In reference to the indigenous chloroplast genes, almost all NUPGs, including *rpl22, atpB, ndhF*, and *rpoC2*, lost their respective coding ability (based on the median of function-retention frequency), whereas more than three-quarters of NUPGs, including *psbl, petN, psaJ*, and *psbN*, maintained their respective coding ability (**Figure 3B**).

**Figure 3.**
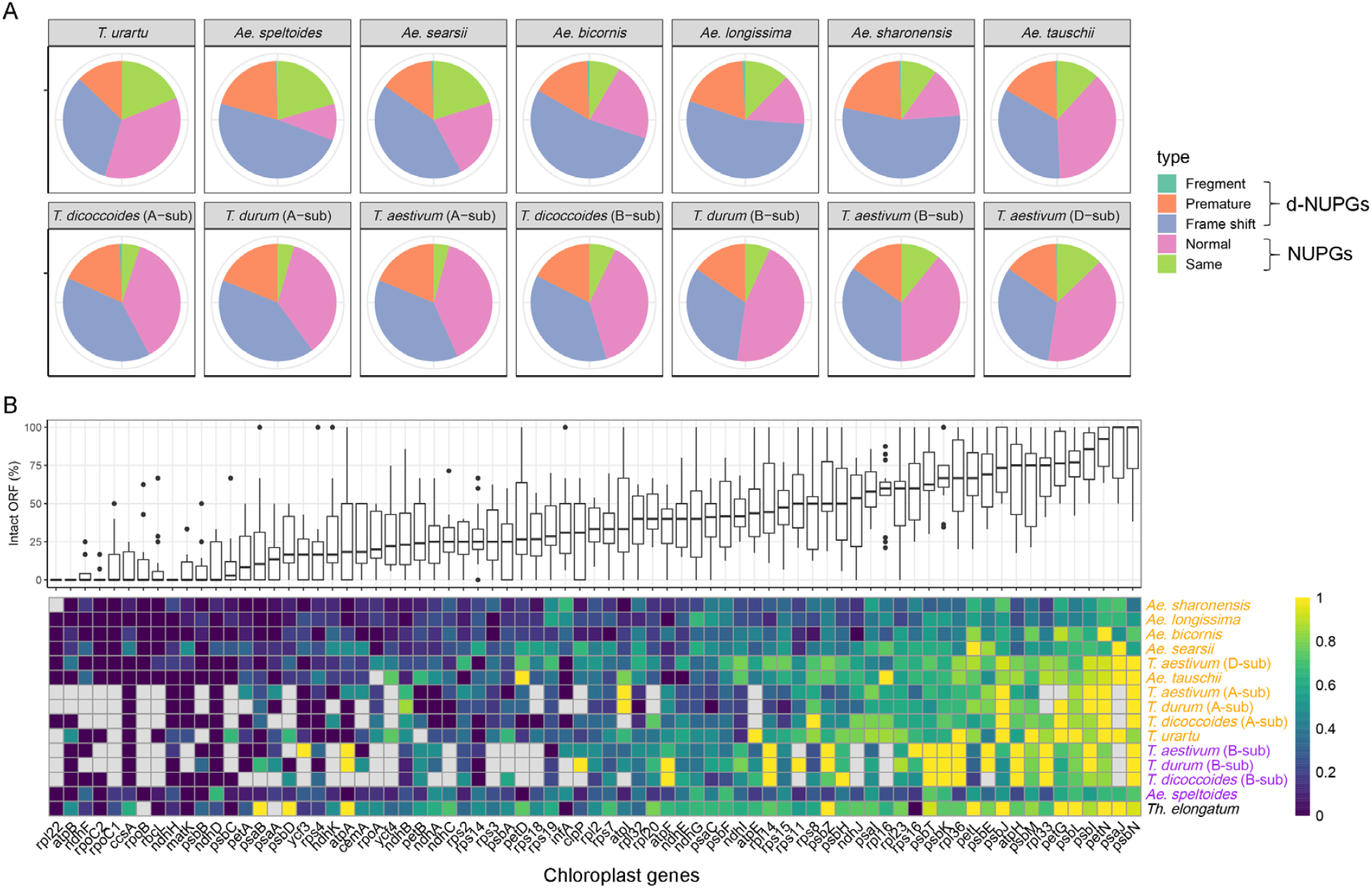
The genetic fate of genes in NUPTs. **(A)** Different genetic fates of genes in NUPTs. Three groups including lost start and terminal codons (fragment), single-nucleotide polymorphism/InDel induced premature (premature), and frameshift were defined as d-NUPT genes (NUPGs) (disruption of open reading frames [ORFs]), whereas those that maintained original (same) and larger than 50% amino acid similarity (normal) with chloroplast gene sequences were defined as NUPGs (maintaining intact ORFs). **(B)** The proportion of intact ORFs (NUPGs) for each chloroplast gene among different genomes/subgenomes. The box plot on the top panel shows the median proportion of each chloroplast gene among the 13 genomes/subgenomes. The heatmap on the bottom panel gives detailed information on each chloroplast gene in each genome/subgenome. The corresponding information on NUMTs is shown in **Figure S5**.

### Possible transcriptional silencing of NUPGs/NUMGs by repressive epigenetic modifications

The eventual coding abilities of NUPGs/NUMGs still depend on their transcriptional status. Accordingly, based on the PacBio SMRT RNA-seq data, we further analyzed whether those NUPGs/NUMGs were transcribed in the nuclear genome of hexaploid wheat (**see Materials and Methods**). We found that 0.04%–2.5% and 0.02%–0.08% of transcripts/isoforms included at least one chloroplast and mitochondrial annotated gene, respectively (**Table 1**). Furthermore, almost all transcripts/isoforms were transcribed from the chloroplast/mitochondrion rather than from NUPGs/NUMGs based on the similarity assessment (**Table 1**), suggesting the global transcriptional silencing of the intact NUPGs/NUMGs after their insertion into the nuclear genome.

**Table 1.** Detection of nuclear plastid DNA/nuclear mitochondrial DNA-derived or organelle-derived transcripts.

| Resource | Tissue | Num.<br>transcripts | Origin of transcripts |  |  | Origin of transcripts |  |  |
| --- | --- | --- | --- | --- | --- | --- | --- | --- |
|  |  |  | (chloroplast genes or NUPGs) |  |  | (mitochondrial genes or NUMGs) |  |  |
|  |  |  | chloroplast | NUPG | ambiguous | mitochondrial | NUM<br>G | ambiguous |
| Kariega | Dusk | 87,246 | 1,291 | 0 | 0 | 22 | 0 | 0 |
| Kariega | Flag leaves | 82,292 | 777 | 0 | 1 | 27 | 0 | 0 |
| Kariega | Root | 80,622 | 32 | 0 | 0 | 35 | 0 | 0 |
| Kariega | Seedling | 47,930 | 687 | 1 | 0 | 11 | 0 | 0 |
| Kariega | Spike | 64,761 | 364 | 0 | 0 | 29 | 0 | 0 |
| Kariega | Grain | 12,474 | 51 | 0 | 0 | 2 | 0 | 0 |
| Zhou8425B | Grain | 51,459 | 49 | 0 | 1 | 8 | 0 | 1 |
| Xiaoyan81 | Grain | 197,709 | 4927 | 3 | 6 | 152 | 0 | 0 |

Considering the importance of epigenetic regulation in transcription, we investigated whether epigenetic regulation can contribute to the foregoing transcriptional silencing. Accordingly, we characterized and compared the epigenetic signal intensities (DNA methylation in the CG, CHG, and CHH context and six histone modifications) among NUPTs/NUMTs, NUPGs/NUMGs, protein-coding genes (PC-genes), transposons (such as *Gypsy, Copia*, and *CACTA* transposons), and their up/downstream flanking regions (+3 kb) (**Figure 4**). Notably, divergent signal patterns were generated for all the aforementioned epigenetic makers between PC-genes and NUPGs/NUMGs (**Figure 4**). Specifically, for DNA methylation, NUPGs/NUMGs with flanking regions were highly methylated; their signal fluctuated across the body, and flanking regions were not as remarkable as that for PC-genes (**Figure 4A**). For the CG and CHG context, the methylation levels of NUMGs were comparable to those of TEs, whereas those of NUMTs, NUPTs, and NUPGs were slightly lower than those of TEs but significantly higher than those of PC-genes (**Figure 4A**). The methylation levels of the CHH context were similar between NUPGs and NUMGs, which were lower than those of NUPTs/NUMTs and TEs (**Figure 4A**). For the euchromatin markers (H3K4me3, H3K27me3, H3K36me3, and H3K9Ac), the signal intensities of PC-genes were significantly higher than those of other genomic features, especially in gene-body regions, whereas NUPGs/NUMGs exhibited the lowest transcriptional activation signal (**Figure 4B**). For the heterochromatin makers, the signal intensities of H3K27me1 in NUPGs and NUMGs were higher than those in PC-genes and lower than those in TEs and NUPTs/NUMTs, whereas those of the NUPGs and NUMGs reached the bottom for H3K9me2 marker (**Figure 4C**). These results suggested that the global transcriptional silencing of NUPGs/NUMGs was mainly attributed to their specific chromatin states, high DNA methylation level, and low level of active epigenetic modifications.

**Figure 4.**
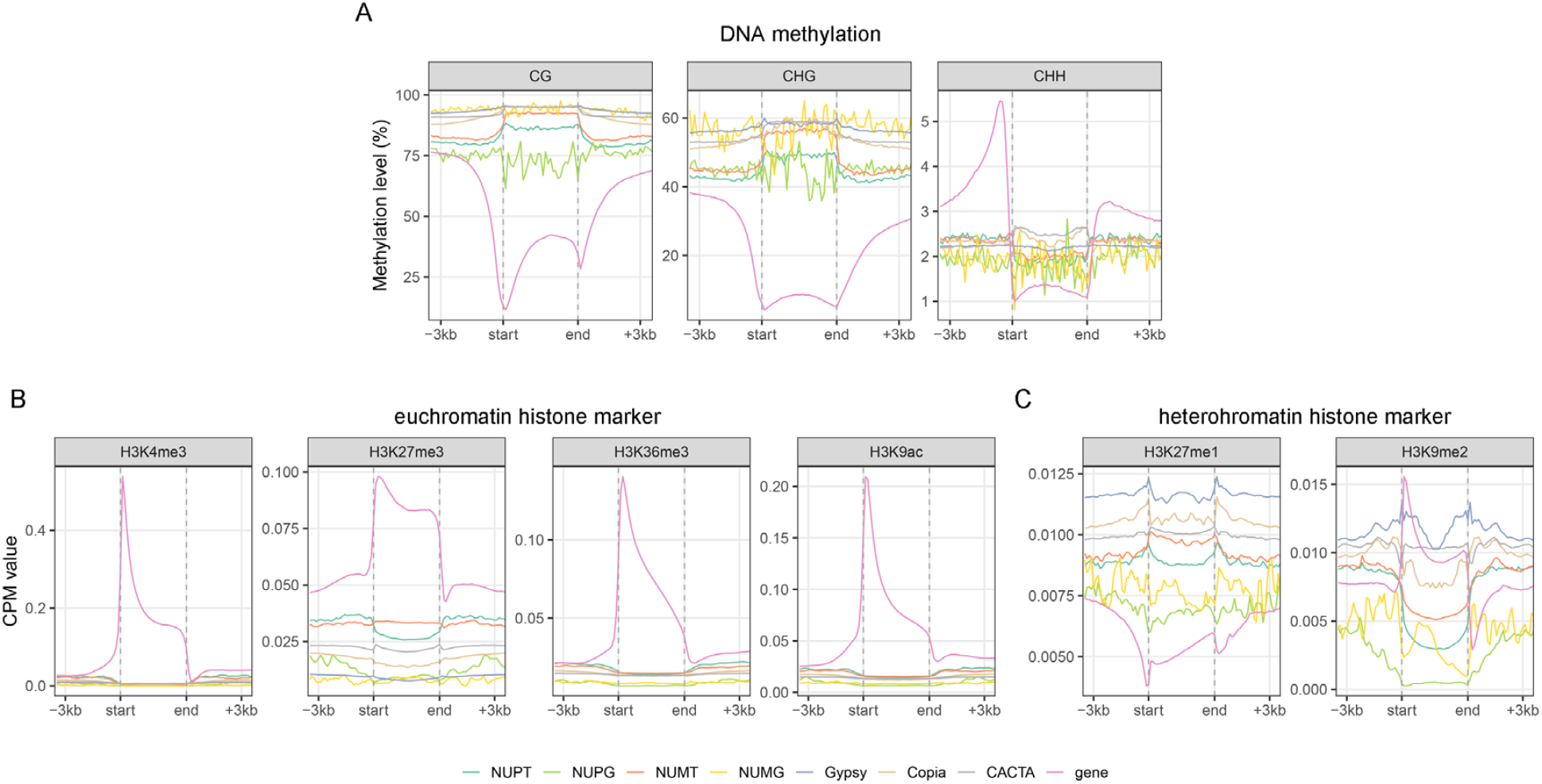
Epigenetic profiling of different genomic features. The read density (for chromatin immunoprecipitation sequencing data, measured as a Counter per Million reads [CPM] value) and methylation signal (for methylome data) of the body and the flanking 3-kb regions of different epigenetic categories including **(A)** DNA methylation (including the CG, CHG, and CHH context), **(B)** representative euchromatin markers (H3K4me3, H4K27me3, H3K36me3, and H3K9ac), and **(C)** heterochromatin markers (H3K27me1 and H3K9me2) were investigated for proteincoding genes, transposable elements (*Gypsy, Copia*, and *CACTA*), NUPTs/NUMTs, and NUPGs/NUMGs, respectively. The regions between dashed lines indicate body region; for protein-coding genes, “start” means the start codon position, whereas “end” means the terminal codon position.

### The gradual relaxation of epigenetic repression in NUPTs/NUMTs

Considering that most of the NUPTs/NUMTs are located in the silent chromatin region, we further investigated the tempo of establishing the current epigenetic status in the alien NUPTs/NUMTs gradually after their insertion. Three possible scenarios were proposed, which included (*i*) gradual heterochromatinization, (*ii*) immediate heterochromatinization maintained over time, and (*iii*) immediate silencing followed by gradually relaxed heterochromatinization. To determine the scenario of the case, we categorized NUPTs/NUMTs into three classes based on their insertion time, which was estimated by their sequence similarity with respective donor segments in the chloroplast/mitochondrial genome: young, similarity ⩾ 98%; intermedium, 94% < similarity < 98%; and old, similarity ⩽ 94%. The epigenetic signal intensities (DNA methylation in the CG, CHG, and CHH context and six histone modifications) of those categorized NUPTs/NUMTs and their up/downstream flanking regions (+3 kb) were characterized and compared as mentioned above.

Regarding DNA methylation, we observed the following: (*i*) the overall hierarchical order of CG DNA methylation levels was “old ≈ intermedium > young” and “old > intermedium ≈ young” for NUPT and NUMT body regions, respectively (**Figure 5A**); (*ii*) the overall hierarchical order of CHG DNA methylation levels was also “old > intermedium ≈ young” for NUMT body regions but was “intermedium > young > old” for NUPT body regions; (*iii*) the hierarchical order “old < intermedium < young” was observed for both of NUPT and NUMT flanking regions in both CG and CHG contexts; (*iv*) the highest DNA methylation level was detected in flanking regions of old NUPTs/NUMTs in the CHH context. For the epigenetic histone modification, except for H3K4me3, the signal intensity of the other four euchromatic markers in body regions of NUPTs and NUMTs increased with their insertion time (**Figure 5B**). Contrary to the active chromatic markers, the two heterochromatic markers existed in the following opposite trend: the signal intensities of old NUPTs and NUMTs were the lowest in both body and flanking regions (**Figure 5C**).

**Figure 5.**
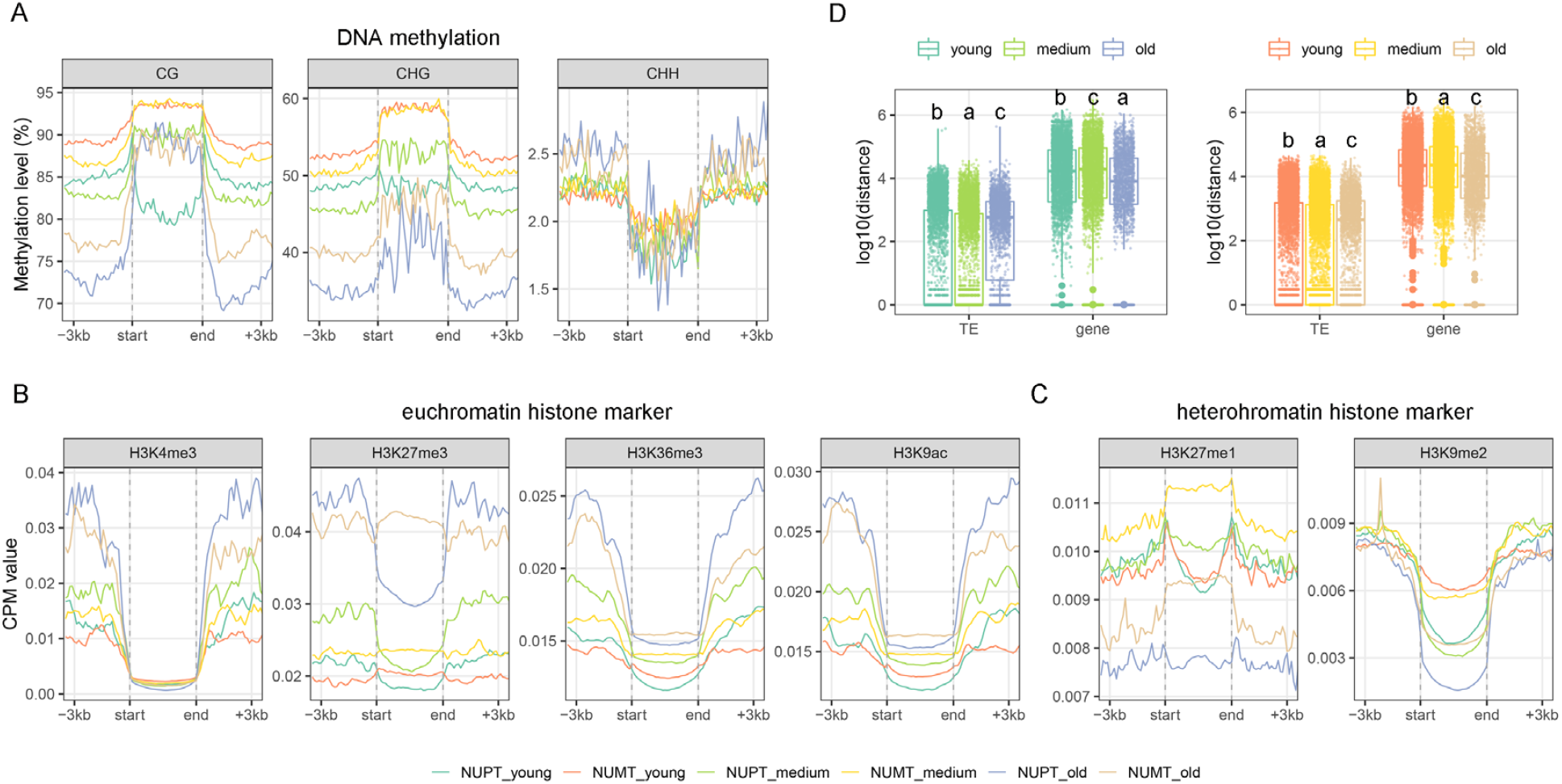
Epigenetic profiling of different types of NUPTs/NUMTs. The read density (for chromatin immunoprecipitation sequencing data and measured as a CPM value) and methylation signal (for methylome data) of the body and the flanking 3-kb regions of different epigenetic categories including **(A)** DNA methylation (including the CG, CHG, and CHH context), **(B)** representative euchromatin markers (H3K4me3, H4K27me3, H3K36me3, and H3K9ac), and **(C)** heterochromatin markers (H3K27me1 and H3K9me2) were investigated for young, medium, and old NUPTs/NUMTs. The regions between dashed lines indicate body regions; for protein-coding genes, “start” means the start codon position, whereas “end” means the terminal codon position. **(D)** Comparisons of the distance between the three NUPT/NUMT groups and adjacent transposable elements and genes. Left panel, NUPTs; right panel, NUMTs. Different letters represent different distances among the three NUPT/NUMT groups (Tukey–Kramer test after Kruskal–Wallis rank sum test, *p* < 2.2e-16).

Interestingly, compared with the young and intermedium NUPTs/NUMTs, the old ones were allocated away from TEs but close to PC-genes (**Figure 5D**; Tukey– Kramer test after Kruskal–Wallis rank sum test, *p* values < 2.2e-16 for both NUPTs and NUMTs). These results suggested that the contextual epigenetic modifications surrounding the nuclear insertion sites underpinned the repressive chromatin status of young (event intermedium) NUPTs/NUMTs, whereas such epigenetic regulation can be gradually relaxed during the course of evolution.

### The gradual accumulation of species-specific NUPTs/NUMTs in the *Triticum*/*Aegilops* complex species

To investigate the evolution of NUPTs/NUMTs at the phylogenic scale, we further strictly identified homologous NUPTs/NUMTs (abbreviated as homo-NUPTs/NUMTs) among the *Triticum*/*Aegilops* complex species and constructed their polymorphism matrix. For two arbitrary NUPTs/NUMTs derived from different genomes, they were classified into an identical homo-NUPT/NUMT group if they had similar body and flanking regions and were located in synteny genomic regions (**Figures S1–S2; see Materials and Methods**). Then, we characterized the dynamic evolutionary history for the homo-NUPT/NUMT group. Taking the NUPT as an example, among seven diploid *Triticum*/*Aegilops* and one outgroup species (*Th. elongatum*), we identified 968 highly confident homo-NUPT groups in which the majority of groups (807; 83.4%) were species-specific (defined as a specific group).

However, only 122 (12.6%) groups were shared by at least two species (defined as a shared group) (**Figure 6A and B**). Following the parsimony criteria, we labeled the dynamic InDels of respective homo-NUPT groups in the phylogenetic speciation tree **(Li, et al. 2022)**. Our findings are as follows: (*i*) the relative insertion frequency of homo-NUPTs increased gradually (on each node) from the ancestral node (3.0; 7.3 MYA) to the present node (35.8; less than 1 MYA) for shared groups; (*ii*) the relative insertion frequencies in the D-lineage species (18.3–25.4) were higher than those in A-(*T. urartu*, 15.0) and B-lineage species (*Ae. speltoides*, 17.3) **(Figure 3C)**, which was consistent with aforementioned NUPT content in D-lineage species (**Figure 1A and B**). Notably, neither species-specific nor shared deletion of NUPTs was detected based on the current 968 homo-NUPT groups, which indicated gradual accumulation during the evolution of the *Triticum*/*Aegilops* complex species. Similarity analysis between NUPTs and respective original chloroplast sequences at each node/tip also supported the accuracy of our current phylogeny-based method **(Figure 6C)**.

**Figure 6.**
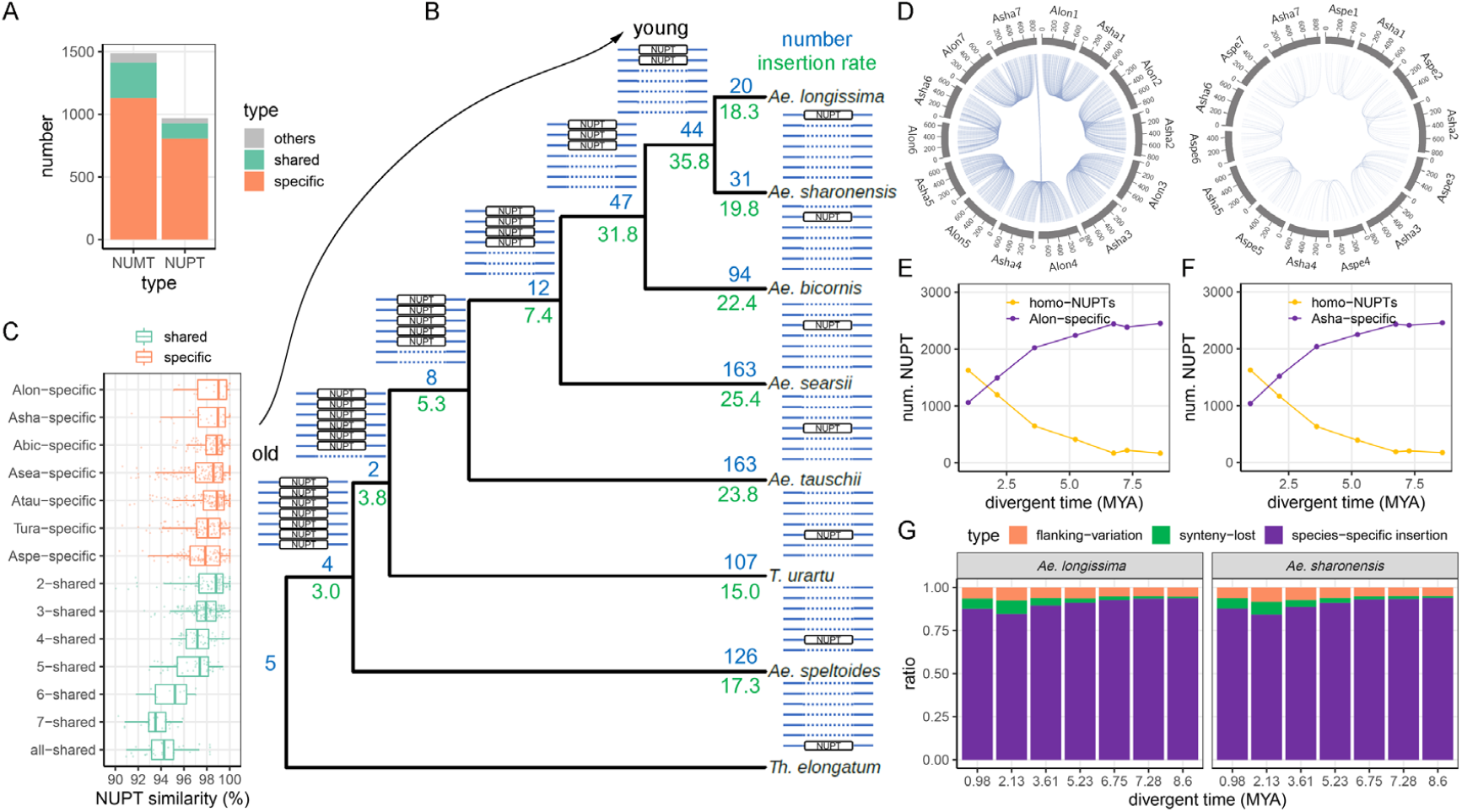
Evolution of NUPTs/NUMTs during species differentiation among diploids from the *Triticum*/*Aegilops* complex species. **(A)** The number of different types of homo-NUPT/NUMT groups. For each homo-NUPT/NUMT group, “shared” means the homo-NUPTs/NUMTs shared by two or more species under each node of the phylogenetic tree (from 2 to 8 species, see **B**), “specific” means only one species including NUPTs/NUMTs, whereas “others” represents the remaining types. **(B)** Phylogeny-based statistics of NUPTs. The ideograms of shared and specific homo-NUPT/NUMT groups are drawn near each node and tip, respectively. Blue and green numbers represent the number and relative insertion ratio (insertion number between two adjacent nodes divided by the evolution time between the corresponding two adjacent nodes) of NUPTs in each node/tip. **(C)** Statistics of NUPT similarity (sequence similarity between NUPTs and corresponding DNA fragments in chloroplast genome sequence) for the shared and species-specific homo-NUPT/NUMT groups. **(D)** Example circus plots of homo-NUPT pairs between *Ae. sharonensis* and *Ae. longissima* (diverged 0.98 MYA) and between *Ae. sharonensis* and *Ae. speltoides* (diverged 7.28 MYA). The numbers indicate the homo-NUPT pairs in each comparison. **(E)–(F)** Change patterns of homo-NUPT pairs and species-specific NUPTs over the divergence time, with *Ae. longissima* **(E)** and *Ae. sharonensis* **(F)** considered as the base (anchor) species, respectively. For each point, the X-axis indicates the divergence time between the base species and one of the rest species. **(G)** The proportion of different types of non-homo-NUPTs in the base (anchor) species when compared with different non-base species. “Flanking variation” means that a given NUPT has a synteny counterpart in the non-base species but the flanking regions are not aligned with each other (the loss of homology at the insertion site). “Synteny lost” means a given NUPT has a counterpart with flanking regions matched but lost synteny relationship. The corresponding information of NUMTs is shown in **Figure S6**. Aspe: *Ae. speltoides*; Tura: *T. urartu*; Atau: *Ae. tauschii*; Asea: *Ae. searsii*; Abic: *Ae. bicornis*; Asha: *Ae. sharonrnsis*; Alon: *Ae. longissima*.

Specifically, older (younger) NUPT groups, which were shared by more (less) species, had lower (higher) sequence similarity. Additionally, the similarity of a given species-specific group was higher than that of its nearest shared group (i.e., the similarity of the *Ae. tauschii*-specific group was less than that of the 5-shared group).

We then performed pairwise homo-NUPT comparisons between *Ae. longissima*/*Ae. sharonensis* (the species at the base of the phylogenetic tree) and each of the rest species. The number of homo-NUPTs decreased from 1,607 (60.2%, *Ae. longissima* vs. *Ae. sharonensis*) to 205 pairs (7.7%, *Ae. longissima* vs. *Th. elongatum*) as the divergent time increased, whereas *Ae. longissima*-specific NUPTs increased from 1,061 to 2,450 (29.8% to 92.3%; **Figure 6D and E**), which was expected. Even though both unaligned flanking regions and non-syntenic NUPTs also contributed to the content of species-specific NUPTs, their proportions were found to be only 5.2%– 17.5% among comparisons across different divergence times. Notably, the proportion of real species-specific insertion increased with divergence time (from 82.5% to 94.8%), whereas the proportion of non-syntenic NUPTs showed an opposite trend (**Figure 6G**). With *Ae. sharonensis* as a comparison anchor, we observed similar results (**Figure 6F and G**). Similar to NUPTs, NUMTs exhibited the species-specific characteristics, and their accumulation gradually increased during the differentiation of the *Triticum/Aegilops* complex diploid species (**Figure S6**).

### Asymmetric ptDNA/mtDNA integration into subgenomes during the evolution of allopolyploid wheat

Contrary to the single-origin nuclear and cytoplasmic genomes in the *Triticum*/*Aegilops* complex diploid species, the uniparental inheritance of maternal organellar genome (B-genome origin) with multi-origin nuclear subgenomes in wild and domesticated allopolyploid *Triticum* species (B- and A-subgenomes in allotetraploid wheat and B-, A-, and D-subgenomes in allohexaploid wheat) allowed us to determine whether ptDNA/mtDNA subgenome integration was asymmetric during the evolution trajectory of allopolyploid wheat (because the real B-genome parent for allotetraploid wheat is still controversial, allotetraploidy process was not considered in our study).

We first compared the profiles of NUPTs/NUMTs in A- and B-subgenomes of wild and domesticated allopolyploid wheat, respectively. As shown in **Figure 7A**, we defined the dynamic index (DI) as the ratio of NUPTs/NUMTs (novel integration into respective subgenomes) that occurred in the stage before compared with after domestication. Accordingly, a significantly higher DI value of a certain subgenome than that of its counterpart represented asymmetric ptDNA/mtDNA integration into different subgenomes in the domestication process. The DI values between A- and B-subgenomes were compared in two different manners, by considering or without considering the corresponding diploid species (**Figure 7B and C;** *T. urartu* and *Ae. speltoides* for A- and B-subgenomes, respectively). When we only considered Chinese Spring as the representative domesticated allohexaploid wheat, the comparison revealed that the DI of NUPTs in the B-subgenome was significantly higher than that in the A-subgenome (0.182 vs. 0.110 if considering diploid species, *p* value = 4,595e-6, **Figure 7B**; 0.145 vs. 0.110 if excluding diploid species, *p* value = 0.0013, **Figure 7C;** Fisher’s exact test). When considering more hexaploid wheat genomes, we found the DI differences mostly supported such subgenomic asymmetry of ptDNA/mtDNA integration into B-subgenome (**Figure 7D and E;** except NUMTs of Norin61 at diploid-including manner, although not all comparisons were statistically significant). Besides foregoing DI comparison, we performed a pairwise comparison of species-specific and -shared NUPTs/NUMTs for paired wild and domesticated allotetraploid wheat species (*T. dicoccoides* vs. *T. durum*), which consistently revealed more species-specific NUPTs/NUMTs after domestication in B-subgenome than in A-subgenome (**Figure 7F**). For hexaploidy, based on the comparison between *T. durum* and numeric *T. aestivum* genomes, we observed increased dominance of polymorphism in B-subgenome (**Figure 7G;** Mann–Whitney U test, *p* value = 1.985e-05 and 8.505e-05 for NUMTs and NUPTs, respectively).

**Figure 7.**
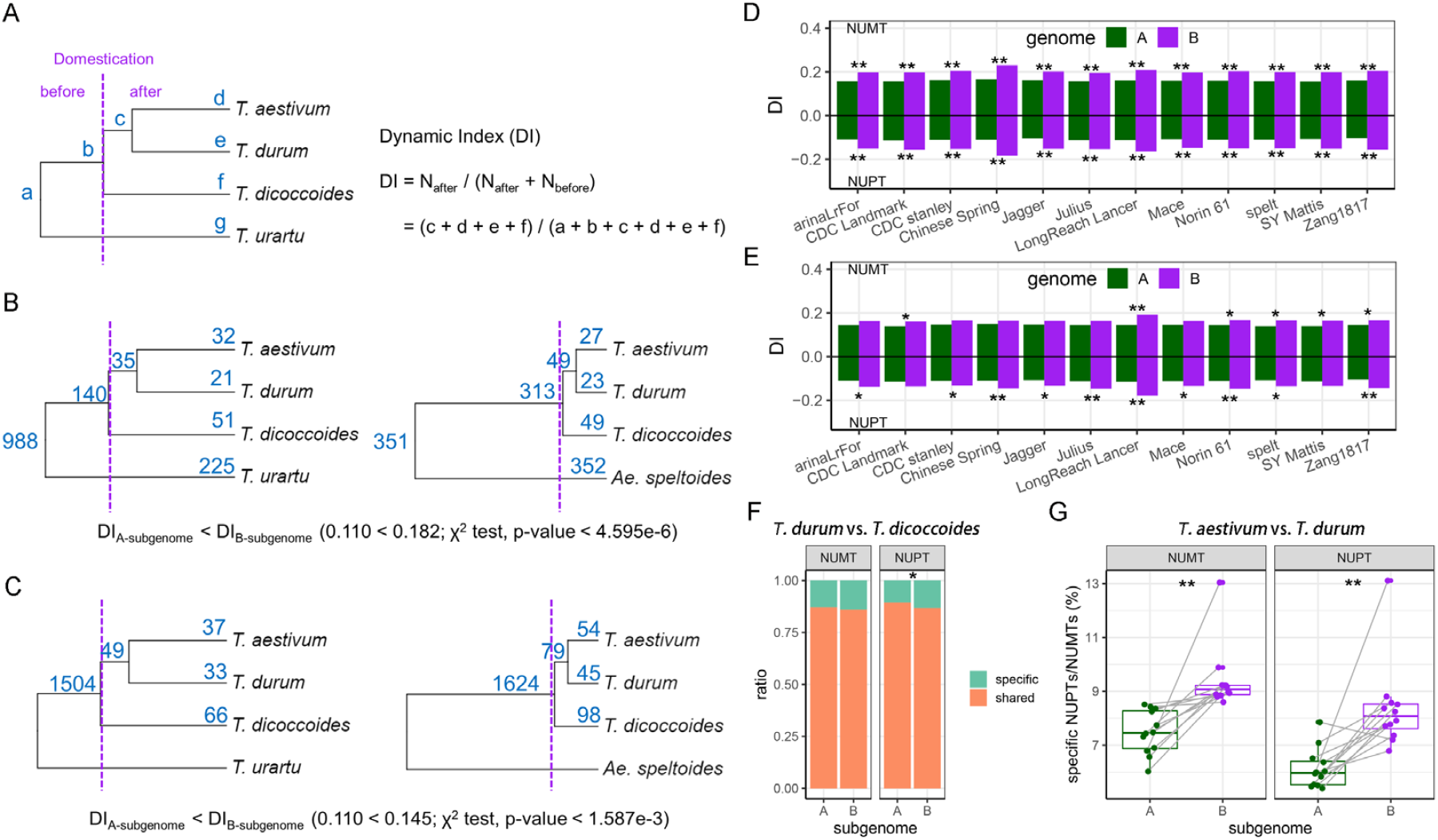
Subgenomic asymmetry of ptDNA/mtDNA integration during tetraploid domestication and allohexaploid processes. **(A)** The schematic diagram for calculating the dynamic index (DI) for NUTPs/NUMTs. The alphabet near each node/tip indicates the number of NUTPs/NUMTs in each evolution node. The dashed line indicates the boundary before and after tetraploid domestication. **(B)–(C)** Comparing the DI of NUPTs between A- and B-subgenomes based on **(B)** diploid-including and **(C)** diploid-excluding manners. Whether the DI in A-subgenome is significantly different from that in B-subgenome is tested using the χ^2^ test. In these two manners, IWGSC RefSeq 1.0 (Chinese Spring) was used as the representative common wheat genome (*T. aestivum*). **(D)–(E)** Comparing the DI of NUPTs/NUMTs between A- and B-subgenomes based on **(D)** diploid-including and **(E)** diploid-excluding manners, using different resources of common wheat genomes. The significant results of the χ^2^ test are shown as ** (*p* < 0.01) and * (*p* < 0.05). **(F)** Statistics of the ratio of the shared to the specific NUPTs/NUMTs based on the number of homo-NUPT/NUMT pairs between *T. dicoccoides* (before domestication) and *T. durum* (after domestication) (χ^2^ test, *p* < 0.05). **(G)** Comparisons of the ratio of specific NUPTs/NUMTs between A- and B-subgenomes based on the number of homo-NUPT/NUMT pairs between *T. durum* (before hexaploidy) and *T. aestivum* (after hexaploidy) using different resources of bread wheat genomes. *p* values were calculated based on the pairwise Mann–Whitney U test (*p* < 0.01).

Finally, to characterize any subgenomic asymmetric accumulation of NUPTs/NUMTs in wheat at the hexaploidy level, we investigated subgenomic polymorphisms of NUPTs/NUMTs in 12 hexaploid wheat genomes from the pangenomic viewpoint. First, the pan-NUPTs were constructed based on homo-NUPTs for each subgenome (**see Materials and Methods and Figure S2**), which revealed their relative abundance as follows: D-subgenome (2,032) > B-subgenome (1,762) > A-subgenome (1,509); the relative abundance of core-NUPTs in the three subgenomes was consistently ranked as follows: D- (1,890) > B- (1,358) > A-subgenome (1,308) (**Figure 8A**). We also calculated the NUPT polymorphism ratio based on the number of core- and pan-NUPTs and found that this ratio was highest in B-subgenome as well (**Figure 8B**; χ^2^ test and post hoc test, *p* value < 0.01). Furthermore, based on the sequenced genomes of wild and domesticated allotetraploid wheat (Zavitan and Svevo, for A- and B-subgenomes) and two *Ae. tauschii* accessions (AL8/78 and AY61 for D-subgenome), we further estimated the gain and loss of NUPTs during the improvement process for each subgenome (**Figure 8C and 8D; Materials and Methods**). As shown in **Figure 8D**, we also found that both gain and loss of NUPTs preferentially occurred in B-subgenome (χ^2^ test and post hoc test, *p* value < 0.01), wherein the gain of NUPTs was significantly higher than the loss of NUPTs (χ^2^ test, *p* value < 0.01). We then performed a pairwise comparison among 12 genomes to characterize the shared ratio of homo-NUPTs (**Figure 8E**), which revealed that the three subgenomes showed significantly different abundance in the shared homo-NUPTs, as follows: B-subgenome < A-subgenome < D-subgenome (**Figure 8F;** Tukey–Kramer test after Kruskal–Wallis rank sum test, *p* value < 0.01). Similar to the results of NUMTs (**Figure S7**), all these results suggested the subgenomic asymmetry of NUPT/NUMT polymorphism during allohexaploidy and the improvement process of wheat.

**Figure 8.**
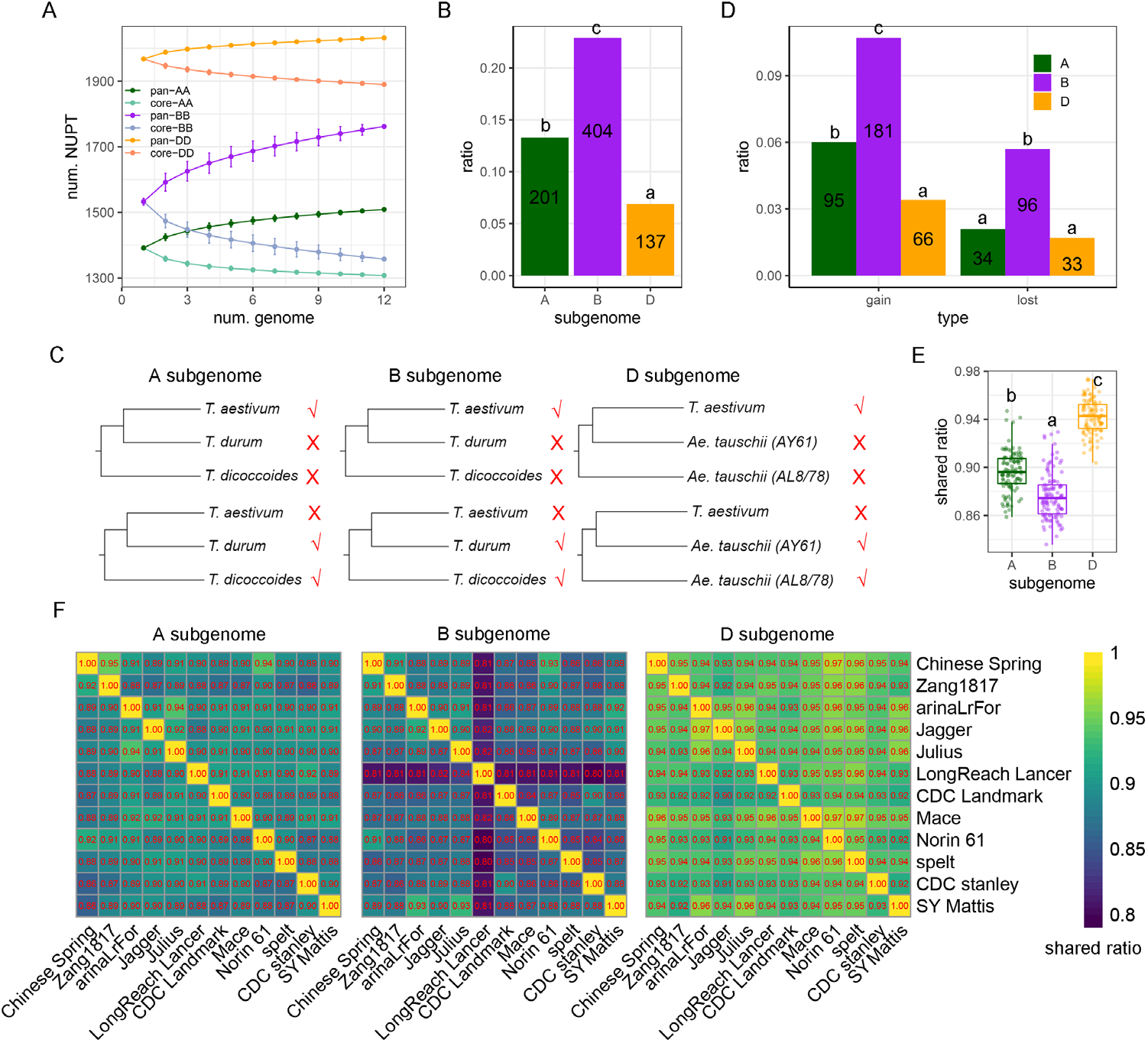
Subgenomic asymmetry of NUPT polymorphism among bread wheat accessions. **(A)** The profiling of pan-NUPTs and core-NUPTs among 12 hexaploid wheat genomes using pan-genome-based analysis according to 1,509 (A-subgenome), 1,752 (B-subgenome), and 2,032 (D-subgenome) homo-NUPT groups. **(B)** Comparisons of the NUPT polymorphism ratio ((N_pan-NUPTs_ − N_core-NUPTs_)/N_pan-NUPTs_) among A-, B-, and D-subgenomes. Alphabets indicate the results of multiple comparisons based on the χ^2^ and post hoc tests. The numbers indicate the difference between the numbers of pan-NUPTs and core-NUPTs. **(C)** Ideogram of gain and loss patterns of NUPTs/nuclear mitochondrial DNAs (NUMTs) among hexaploid genomes based on two outgroup genomes for each of the three subgenomes. Taking A-subgenome as an example: the outgroup genomes are A-subgenomes of *T. dicoccoides* and *T. durum*. For each homo-NUPT group, if NUPTs did not occur in both outgroup genomes but existed in hexaploidy genomes (at least one of 12 genomes), it is treated as a gain group; if NUPTs occurred in both outgroup genomes but did not exist in at least one hexaploidy genome, it is treated as a lost group. **(D)** The proportion of gain and loss homo-NUPT groups among the three subgenomes in hexaploid wheat species. Alphabets indicate the results of multiple comparisons based on the χ^2^ and post hoc tests. The number of gain and loss groups is also shown. **(E)** Pairwise comparisons of homo-NUPT pairs among 12 genomes of the three subgenomes. The number in each cell represents the proportion of homo-NUPTs to the total number of NUPTs for each comparison. **(F)** Summary of homo-NUPT ratios among the three subgenomes based on 12 hexaploid genome datasets. Alphabets indicate the results of multiple comparisons based on the Kruskal–Wallis rank sum test and Tukey–Kramer test (*p* < 2.2e-16). The corresponding information on NUMTs is shown in **Figure S7**.

## Discussion

Integration of organellar DNA (both mitochondrial and/or chloroplast DNA) into the nuclear genome has been identified in many eukaryotes from fungi and plants to mammals, which affects the genome structure and genetic diversity and further promotes evolution **(Leister 2005; Kleine, et al. 2009; Sloan, et al. 2018)**.

Nevertheless, the transcriptional expression and epigenetic state of organelle genes inside NUPTs/NUMTs and the evolutionary dynamics of NUPTs/NUMTs at the phylogenic scale are poorly explored. Accordingly, in the *Triticum*/*Aegilops* complex species with abundant NUPTs/NUMTs and distinct evolutionary trajectories **(Marcussen, et al. 2014; Glémin, et al. 2019; Zhang, et al. 2020)**, we determined the genetic mutation, transcriptional expression, and epigenetic status of NUPTs/NUMTs and their phylogenomic and pan-genomic insertion characteristics during diploid speciation, polyploidization, and domestication.

### Transcriptional silencing and epigenetic control of NUPGs/NUMGs

A previous study showed that organelle-derived nuclear genes are always inactivated, lose their original function, and are pseudogenized **(Kleine, et al. 2009)**. Consistently, we found most NUPGs/NUMGs identified in the *Triticum*/*Aegilops* complex species were pseudogenized after the accumulation of genetic mutations that cause ORF interruption. However, the NUPGs/NUMGs maintaining intact ORFs facilitated the determination of the potential epigenetic regulation underlying respective inactivation. Accordingly, our methylome and ChIP-seq analyses showed that ORF-intact NUPGs/NUMGs were significantly distinct from endogenous nuclear genes but similar to TEs in terms of their epigenetic modifications (**Figure 4**). Consistent with previous findings that showed that NUPTs/NUMTs and TEs might share a similar homology-dependent DNA methylation mechanism to maintain nuclear genome stability **(Maumus and Quesneville 2014; Yoshida, et al. 2019)**, we confirmed that NUPGs/NUMGs did not possess the epigenetic properties of actively transcribed genes. Furthermore, the full-length transcriptomic analysis confirmed that almost all NUPGs/NUMGs are transcriptionally silent compared with their counterparts in the organelles (**Table 1**). Accordingly, as a novel input to the fate of NUPGs/NUMGs, ORF-intact NUPGs/NUMGs can still be transcriptionally silenced under epigenetic control.

Another well-known fate of NUPGs/NUMGs is functional maintenance in encoding proteins targeting back to the original endosymbionts, such as proteobacteria-like and cyanobacteria-like prokaryotes, which involves exemplary *rbcS* encoding subunits of the chloroplast RuBisco complex and *cytochrome c* encoding subunits of the mitochondrial enzyme complex of oxidative phosphorylation **(Blier, et al. 2001; Rand, et al. 2004; Andersson and Backlund 2008)**. How those ancient NUPGs/NUMGs escaped from foregoing transcriptional silencing is an intriguing question. Given that certain NUPGs/NUMGs identified in rice species were integrated into euchromatic regions **(Wang and Timmis 2013)**, the NUPGs/NUMGs generating foregoing functional genes possibly integrated into the euchromatic regions with a low load of epigenetic silencing within their eukaryotic ancestors.More epigenetic and evolutionary data with the basal eukaryotic species are required to test this hypothesis.

### Species-specific and continuous insertion mechanisms for NUPTs/NUMTs

Previous studies have compared the genomic composition and characteristics of NUPTs/NUMTs within various species **(Michalovová, et al. 2013; Zhao, et al. 2019)**. However, as they rarely distinguish shared- and species-specific NUPTs/NUMTs, making characterization of the dynamics of NUPTs/NUMTs in the evolution of closely related species is difficult **(Liang, et al. 2018)**. In the present study, we used a phylogenomic-based method to identify homo- and species-specific NUPT/NUMT groups among diploid *Triticum*/*Aegilops* complex species for the robust investigation of NUPTs/NUMTs at the evolutionary scale. The results suggested that the species-specific insertion of NUPTs/NUMTs, having significantly higher abundance than homo-NUPTs/NUMTs, mainly contributed to NUPT/NUMT polymorphism within the last ∼7 million years. Moreover, by using the phylogenomic method, we also estimated and compared the relative insertion rates of those NUPTs/NUMTs **(Richly and Leister 2004; Leister 2005)**. Interestingly, we observed a gradual increase in the relative insertion frequency of homo-NUPTs from the ancestral node to the present node (**Figure 6B and S6B**). The relevant results showed that the insertion rate was related to the number of NUPTs/NUMTs (for NUPTs, D-lineage species > *Ae. speltoides* > *T. urartu*; for NUMTs, D-lineage species > *T. urartu* > *Ae. speltoides*), indicating that the evolution of NUPTs/NUMTs occurred gradually in diploid *Triticum*/*Aegilops* complex species.

Notably, the relative abundance of homo-NUPTs/NUMTs was low in the respective species. Such a phenomenon could be explained by two possible events, the involvement of the loss of homology at the insertion site and limited synteny. TE insertion, rapid mutation, and deletion could all cause the former homology loss of the insertion site; considering the non-synteny feature of TEs in the three subgenomes of common wheat **(Wicker, et al. 2018)**, the majority of ptDNA/mtDNA inserted near TEs lacked collinearity between the highly diverged species.

### Integration asymmetry of ptDNA/mtDNA among subgenomes during the evolution trajectory of allopolyploid wheat

Besides empirical evidence for subgenomic dominance (bias in gene retention, gene diversity, TE dynamics, and epigenetic modifications) in allopolyploids **(Pont and Salse 2017; Li, et al. 2021; Levy and Feldman 2022)**, we observed and defined the novel subgenomic dominance regarding the asymmetric integration of ptDNA/mtDNA into different subgenomes (majorly biased integration into the B-subgenome). As both A- and D-subgenomes were less fractionated and B-subgenome was more fractionated during allopolyploid wheat evolution **(Pont, et al. 2013; El Baidouri, et al. 2017; Pont and Salse 2017)**, the more plastic subgenome increased NUPT/NUMT polymorphism. We assume that a subgenome with more TE dynamics may generate more TE-related DSBs, especially during meiosis, which eventually enhances the dynamics of NUPTs/NUMTs. This assumption is based on the following lines of evidence: (*i*) the molecular basis for the integration of ptDNA/mtDNA necessitates nuclear DSBs **(Hazkani-Covo, et al. 2010)**, (*ii*) TE transportation generally produces DSBs **(Gorbunova and Levy 1999; Gasior, et al. 2006; Hedges and Deininger 2007)**, and (*iii*) ptDNA/mtDNA preferably inserted in TE-related regions in the present *Triticum*/*Aegilops* case. Moreover, B-subgenome, which is the largest subgenome and contains more abundant TEs in the wheat genome **(Consortium, et al. 2018)**, can carry a stronger genetic load, including ptDNA/mtDNA insertions. Notably, a comprehensive introgression into the B-subgenome of allopolyploid wheat **(Walkowiak, et al. 2020; Zhou, et al. 2020; Wang, et al. 2022)** may also be a potential contributor to enhance the genomic diversity and further NUPT/NUMT polymorphism. Additional molecular and evolutionary evidence is required to validate these speculations in the future.

Taken together, the present systematic analyses of the whole-genome atlas of NUPTs/NUMTs in the *Triticum*/*Aegilops* complex species reveal their repressed epigenetic status, species-specificity, gradual accumulation, and asymmetric subgenome integration in allopolyploid species. The study provides new insights into the evolution of nuclear organellar DNAs in plants.

## Materials and Methods

### Sequence resources of nuclear, chloroplast and mitochondria

All genome sequence resources of nuclear, chloroplast and mitochondria were obtained from previous publications and/or NCBI website **(https://www.ncbi.nlm.nih.gov/)**. Nuclear genome sequences included *Thinopyrum elongatum, Triticum urartu* **(Ling, et al. 2018)**, five *Aegilops tauschii* accessions (AL8/78, AY17, AY61, T093 and XJ02) **(Luo, et al. 2017; Zhou, et al. 2021)**, five Sitopsis species (*Ae. speltoides, Ae. searsii, Ae. bicornis, Ae. sharonensis* and *Ae. longissima*) **(Li, et al. 2022)**, *T. dicoccoides* **(Avni, et al. 2017)**, *T. durum* **(Maccaferri, et al. 2019)**, *T. aestivum* of semi-wild Zang1817 **(Guo, et al. 2020)** and extra 11 accessions (Chinese Spring, Arina*LrFor*, Jagger, Julius, LongReach Lancer, CDC Landmark, Mace, Norin61, Spelta, CDC Stanley and SY Mattis) **(Consortium, et al. 2018; Walkowiak, et al. 2020)**. Chloroplast genome sequences were obtained from NCBI website, including *Th. elongatum* (NC_043841), *T. urartu* (MG958555), *Ae. tauschii* (MG958544), *Ae. speltoides* (MG958553), *Ae. searsii* (NC_024815), *Ae. bicornis* (NC_024831), *Ae. sharonensis* (NC_024815), *Ae. longissima* (MG958549), *T. dicoccoides* (MG958552), *T. durum* (MG958545) and *T. aestivum* (MG958554). Mitochondria genome sequence of *T. aestivum* (NC_036024) was also obtained from NCBI. For polyploid nuclear genomes, they were split to different subgenome sequences to separate database files for further identification of NUPTs/NUMTs.

### Identification of NUPTs/NUMTs and intra-genomic duplication NUPTs/NUMTs

BLAST based method was used to identify NUPTs/NUMTs. For NUPTs, all query chloroplast sequences were aligned to corresponding or related (for example, for identification of NUPTs in *T. aestivum* A-subgenome, the chloroplast genome of *T. urartu* was treated as query sequence) genomes/subgenomes using Blastn. While for NUMTs, the mitochondria genome of *T. aestivum* was aligned to each genomes/subgenomes using Blastn. The parameters of Blastn were set as “-evalue 1e-10 -dust no -penalty -2 -word_size 9”. To obtain high confident NUPTs/NUMTs, the raw NUPT/NUMT hits were further filtered based on the following criteria: (i) the length of Blast hit is larger than 100bp; (ii) the similarity of Blast hit is larger than 90%; (iii) if the distance between adjacent NUPTs/NUMTs is less than 1kb, they were merged into one NUPT/NUMT. The annotation of the genomic regions for NUPTs/NUMTs was performed using ChIPseeker **(Yu, et al. 2015)**. The nearest genomic features (such as gene and different types of TEs) of NUPTs/NUMTs were determined using Bedtools **(https://bedtools.readthedocs.io/en/latest/index.html)**.

The intra-genomic duplication of NUPTs/NUMTs after their insertion in nuclear genome were identified based on previous methods with modification **(Liang, et al. 2018)**: (i) all vs. all Blastn of NUPTs/NUMTs in each genome/subgenome was performed for checking candidate NUPT/NUMT pairs; (ii) for each duplicated NUPT/NUMT pair, the 5’ and 3’ flanking regions of 500bp length were extracted for further pairwise alignment using Blastn. The candidate NUPT/NUMT pair was retained if both of the flanking regions were well aligned; (iii) MCscanX **(Wang, Tang, et al. 2012)** was performed to classify maintained candidate NUPT/NUMT pairs to duplication categories, including dispersed, tandem, proximal and segmental duplication NUPTs/NUMTs.

### Detection of homo-NUPTs/NUMTs groups among genomes/subgenomes

24,239 conserved genomic regions (CGRs) among eight diploid genomes (including seven *Triticum/ Aegilops* species and *Th. elongatum*) were constructed based on our previous pipeline **(Li, et al. 2022)**. In brief: first, the genic and flanking 20kb regions of all diploid genomes were aligned to the B-subgenome sequence of IWGSC RefSeq 1.0 (backbone sequence) using the nucmer module of MUMmer v3.9 **(Kurtz, et al. 2004)** with the parameters --mum -c 90 -l 40. Second, the best one-to-one query-reference alignments for each diploid genome were obtained based on delta-filter and show-coords module. Then, Bedtools *intersect* module was used to identify original CGRs among all species based on the alignment regions on the backbone sequence. Finally, the original CGRs from backbone genome were re-aligned to each diploid genome sequence for detection of final CGRs through minimap2 (v2.17) **(Li 2018)**. According to above 24,239 CGR markers, we also constructed 22,763, 23,287 and 22,531 CGR markers for A-, B- and D-subgenomes of 12 *T. aestivum* genomes, respectively. The CGR marker showed remarkable syntenic relationships among different genomes and therefore provided robust anchors for further homo-NUPTs/NUMTs detection. The CGR markers were ranked in each genome/subgenome.

The method for detection of homo-NUPTs/NUMTs between a given pair of genome/subgenomes was mimic to intra-genomic duplication NUPTs/NUMTs. A pair of NUPTs/NUMTs from two different genome/subgenomes were defined as homo-NUPT/NUMT if: (i) their NUPT/NUMT bodies were well aligned; (ii) their NUPT/NUMT flanking regions (500 bp) were well aligned; (iii) they were located in synteny regions, i.e., the rank difference between the NUPT/NUMT-nearest CGR markers was less than 20. Except homo-NUPTs/NUMTs, the remaining NUPTs/NUMTs in each genome/subgenome were defined as genome/subgenome-specific NUPTs/NUMTs. For given genome/subgenome-specific NUPT/NUMT, if its flanking regions have syntenic locus in the other genome/subgenome, we defined such locus as NUPT/NUPT-related homologous locus (NHL). The homo-NUPT/NUMT pairs and NUPT/NUMT-NHL pairs were used for further identification of homo-NUPT/NUMT groups.

For the eight diploid genomes, we performed all possible pairwise comparisons and obtained 28 combinations of homo-NUPT/NUMT pairs and NUPT/NUMT-NHL pairs. We used the Python model *networkx* to concatenate all related homo-NUPT/NUMT pairs and NUPT/NUMT-NHL pairs and generate candidate homo-NUPT/NUMT groups among eight species. If a given homo-NUPT/NUMT group satisfied (i) including exact eight members, with each species providing one member and (ii) possessing only two types of members namely NUPT/NUMT and NHL, it will be maintained as a diploid-level homo-NUPT/NUMT group for further analysis. For 12 hexaploid wheat genomes, we performed same analysis to obtain A-, B- and D-subgenome-level homo-NUPT/NUMT groups (also defined as A-, B- and D-subgenome pan-NUPTs/NUMTs), respectively. The ideogram of the pipeline on identification of homo-NUPT/NUMT groups were shown in **Figure S2**.

### Variant calling for NUPTs/NUMTs and coding ability checking for organelle-derived genes

For each genome/subgenome, NUPTs/NUMTs were first aligned to corresponding organelle genome sequences and generated related Bam files using Minimap2.

Second, Samtools *mpileup* module (**https://www.samtool.org**) was performed to produce the pileup files which including the base information for each nucleotide site of organelle genome sequences. Custom Python scripts was used to parse the pileup file and obtain the variant information (including SNP and InDels) and insertion times (depth) for each non-overlapping 1kb nucleotide region.

For NUPTs/NUMTs which covered the complete gene body regions of genes in organelle genome sequences, we aligned them to corresponding coding sequences of organelle genes by MAFFT (https://mafft.cbrc.jp/alignment/software/). Custom Python script was used to search variant sites (including SNPs and InDels) occurred in CDS regions which changed the coding ability of organelle genes. Six possible destinies of organelle-derived genes might be happened: (i) identical to original organelle gene (same); (ii) SNP/InDel introduced variations that cause amino acid changes, but ORF region maintained and have more than 50% sequence similarity to the source gene (normal); (iii) SNP/InDel introduced variations that cause amino acid changes, but ORF region maintained and have less than 50% sequence similarity to the source gene (new); (iv) SNP/InDel-induced premature; (*v*) SNP/InDel-induced loss of initial and stop codons (fragment) and (*vi*) frame shift (the sequence length of alignment region is not multiple of three). The first two types maintained the intact ORF of source organelle genes were defined as NUPT/NUMT-related genes (NUPGs/NUMGs), whereas the last three types with disrupted ORF were defined as d-NUPGs/NUMGs.

### RNA-seq data analysis

Considering the powerful potency for identification of full-length RNA transcripts, the PacBio SMRT RNA-seq data were used to determine whether an expressed gene is derived from chloroplast/mitochondria or nuclei. Previous published full-length transcript datasets of hexaploidy wheat **(Dong, et al. 2015; Wei, et al. 2019; Athiyannan, et al. 2022)**, which assembled based on long sequencing reads from Pacbio SMRT platform, were download from NCBI SRA and GEO database (including ERR6022024, ERR6022025, ERR6022026, ERR6022027, ERR6022028, ERR6022029, SRR3018829 and GSE118474). The transcript datasets were aligned to the reference genome and transcriptome of both nuclear (IWGSC reference genome V1.0) and organelle (MG958554 for chloroplast and NC_036024 for mitochondria) of heaxploid wheat using Minimap2. If the editing distance (based on the number of SNPs and InDels) of a given transcript to the NUPG/NUMG is less than that to the organelle gene, such transcript is inferred as NUPG/NUMG -derived, otherwise it is inferred as chloroplast/mitochondria-derived transcript/isoform.

### Methylome and ChIP-seq data analysis

The hexaploid wheat methylome data (SRR6792673, SRR6792681, SRR6792684, SRR6792687, SRR6792688 and SRR6792689) and ChIP-seq data including ASY1 (ERR464976), DMC1 (ERR4649761), H3K4me3 (ERR4649763), H3K27me3 (SRR10300747), H3K27me3 (SRR6350666), H3K9ac (SRR6350667), H3K36me3 (SRR6350670), H3K27me1 (ERR4649762), H3K9me2 (ERR4649764) were download from NCBI SRA database **(Consortium, et al. 2018; Tock, et al. 2021)**. A combined reference genome data was constructed through merging IWGSC RefSeq 1.0, chloroplast (MG958554) and mitochondria (NC_036024) genome sequences for further short reads alignment.

For methylome data analysis, after filtering out adaptors and low-quality data by Trimmomatic (Bolger *et al*., 2014), the bisulfite-treated short reads were first aligned to the combined reference genome using bismark **(Krueger and Andrews 2011)** with default parameters. Second, *bismark_methylation_extractor* and *bismark2bedGraph* modules were performed to generate bedGraph files for CG, CHG and CHH genomic context. Then, bedGraph files were converted to bigWig files using *bedGraphToBigWig* script (https://www.encodeproject.org/software/bedgraphtobigwig/). Finally, deepTools **(Ramírez, et al. 2014)** *computeMatrix* module was used to calculate the methylation level of different genomic features (including genes, TEs, NUPTs/NUMTs and NUPGs/NUMGs) and flanking regions (3000 bp) in CG, CHG and CHH genomic context.

For ChIP-seq data analysis, after filtering out adaptors and low-quality data by Trimmomatic, all short reads datasets were first aligned to the combined reference genome using Bowtie2 **(Langmead and Salzberg 2012)** with default parameters.Second, the aligned reads which satisfied (i) proper pair (ii) MAPQ large than 2 and (iii) less than 6 mismatches were maintained. The filtered Bam files were converted to bigwig file using deepTools *bamCoverage* module. Finally, deepTools *computeMatrix* module was used to calculate the reads density of different genomic features (including genes, TEs and NUPTs/NUMTs and NUPGs/NUMGs) and flanking regions (3000 bp).

## Acknowledgements

This work was supported by the National Natural Science Foundation of China (NSFC #31970238), the China Postdoctoral Science Foundation (#2021M700749), the Young Scientific and Technological Talents Supporting Project of Jilin Province (#QT202119) and the Fundamental Research Fund for Central Universities.

## Author Contributions

Z.B.Z., B.L., and L.G. designed the research. Z.B.Z., B.L., and L.G. performed the research. B.W. and Y.Q.M. collected and preprocessed all sequencing data, Z.B.Z., J.Z., J.Z.L., J.T.Y., N.L., and T.Y.W. analyzed data. Z.B.Z., H.Y.W., B.L., and L.G. wrote the manuscript.

## Competing interest statement

The authors declare no competing interests.

## Notes

### Competing Interest Statement

The authors have declared no competing interest.

