## supplementary figures for "Evolutionary trajectory of organelle-derived nuclear DNAs in the *Triticum/Aegilops* complex species"

Figure S1-S7

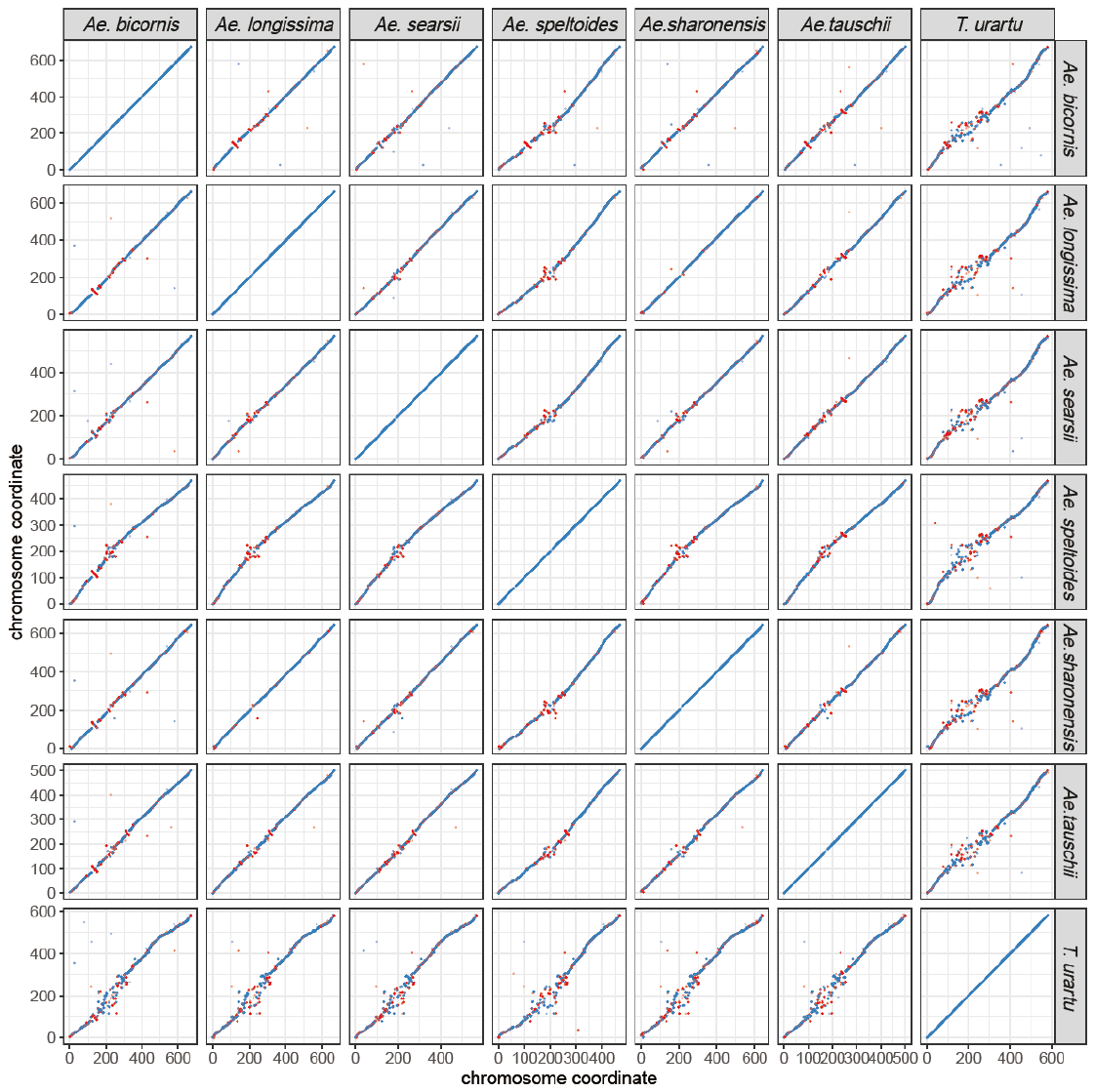

**Figure S1. Dot plots of conserved genomic regions (CGRs) among seven diploid species in the *Triticum/Aegilops* complex.** Identification of CGRs is detailed in the Materials and Methods section. Only CGRs of chromosome 1 are shown.

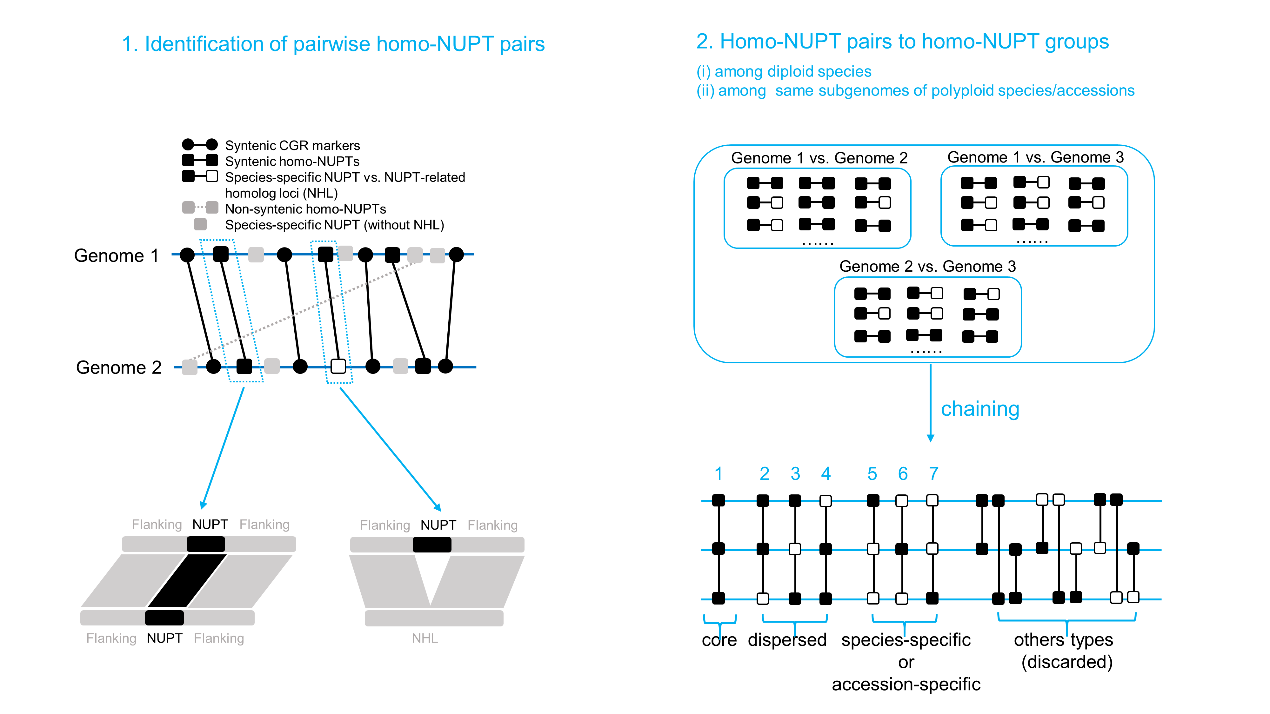

**Figure S2. Flow chart for the identification of homo-NUPT/NUMT groups.** The concept of homo-NUPT/NUMT groups is used for both (*i*) eight diploid species (phylogeny level) and (*ii*) 12 hexaploid accessions as well as two tetraploid species (including A, B, and D subgenomes, pan-genome level). Only NUPT-NUPT (NUMT-NUMT) and NUPT-NHL (NUMT-NHL) pairs (NHL means NUPT/NUMT-related homolog loci) between each pair of genome/subgenome are used for the construction of homo-NUPT/NUMT groups among (*i*) diploid species and (*ii*) same subgenomes of polyploidy species/accessions. Homo-NUPT/NUMT groups lack of NUPT/NUMT and NHL information are discarded for further analysis. The seven example homo-NUPT/NUMT groups which satisfy the subsequent analysis were shown.

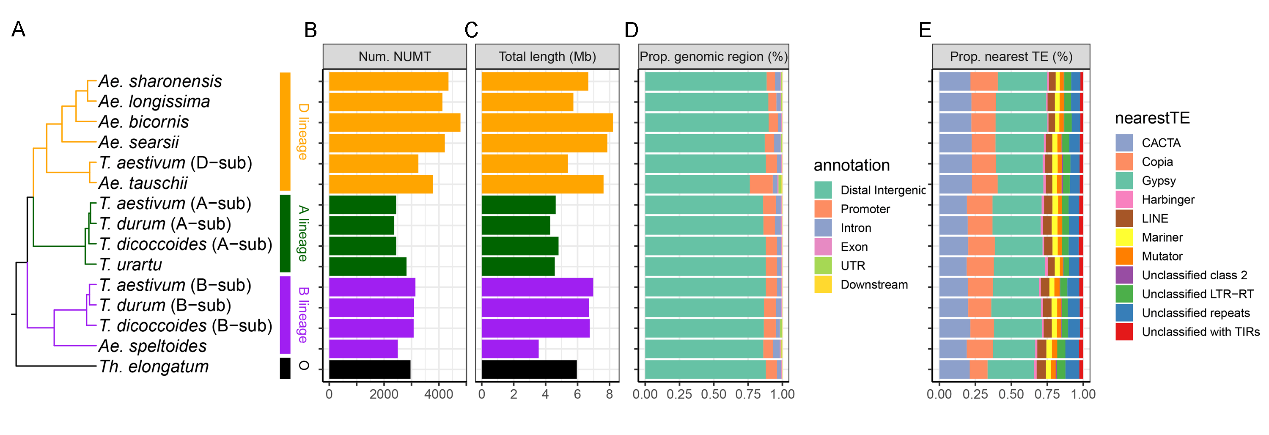

**Figure S3. Genomic landscape of NUMTs in *Triticum/Aegilops* complex species. (A)** Phylogeny tree topology of representative *Triticum/Aegilops* complex genomes、subgenomes. Whole genome statistics of NUPTs including **(B)** numbers **(C)** total length **(D)** the proportion of distribution on different genomic features and **(E)** the proportion of nearest transposon types. The corresponding genomic characteristics of NUPTs are shown in **Figure 1.**

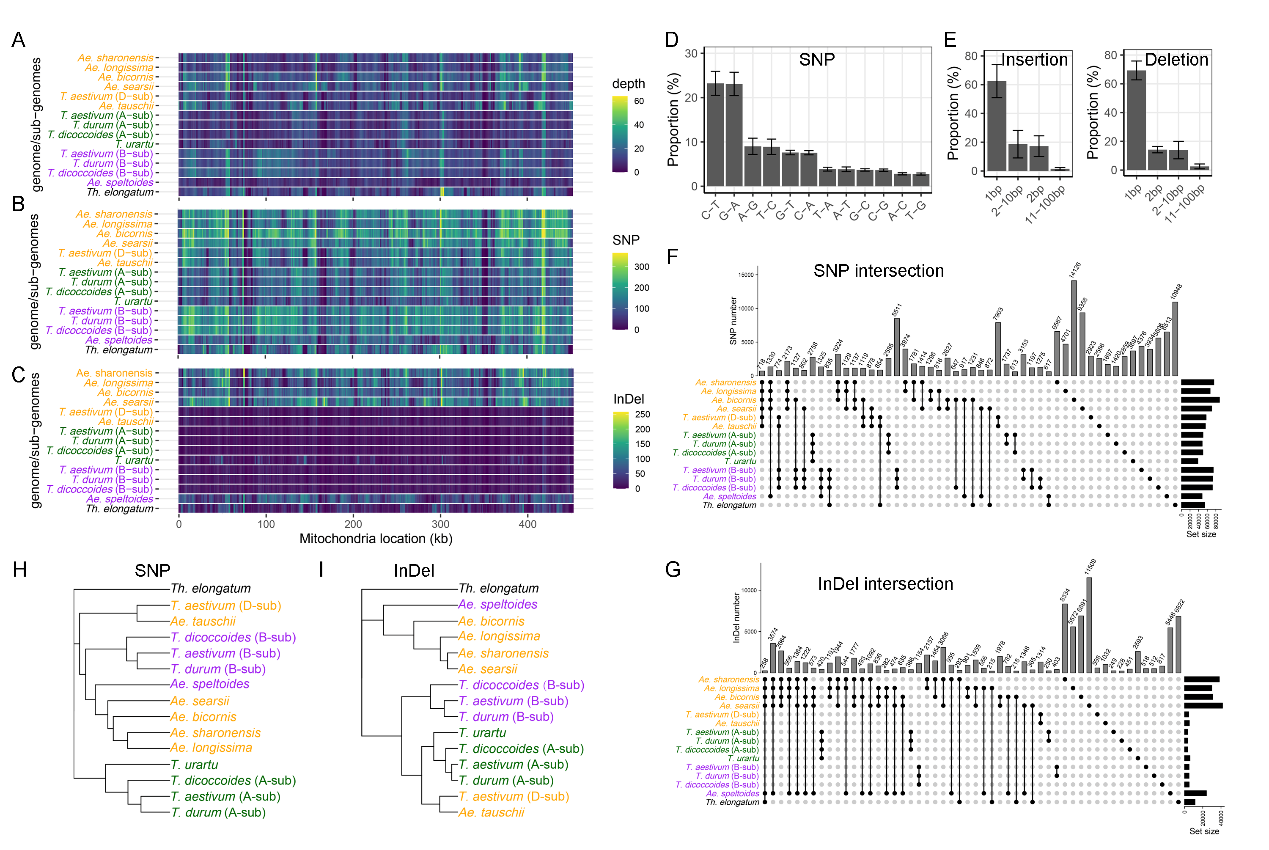

**Figure S4. Characteristics of genetic variations in NUMTs (compared with mitochondria genome).** Density feature of NUMTs in different regions of chloroplast genome based on non-overlapping 1kb windows, including **(A)** insertion frequency (depth), **(B)** SNP density and **(C)** InDel density. NUMTs were first aligned to chloroplast genomes and then calculated for each feature. **(D)** The proportion of different types of SNPs. The error bars indicate the standard deviation (SD) among different genomes/subgenomes. **(E)** The proportion of different types of insertions and deletions. **(F) – (G)** The UpSet plot based on the intersection matrix of SNPs **(F)** and InDels **(G)** in each variation site among genomes/subgenomes. **(H) – (I)** The NJ tree topology based on the intersection matrix of SNPs **(H)** and InDels **(I)** used in **(F)** and **(G)**. The corresponding information of NUPTs is shown in **Figure 2**.

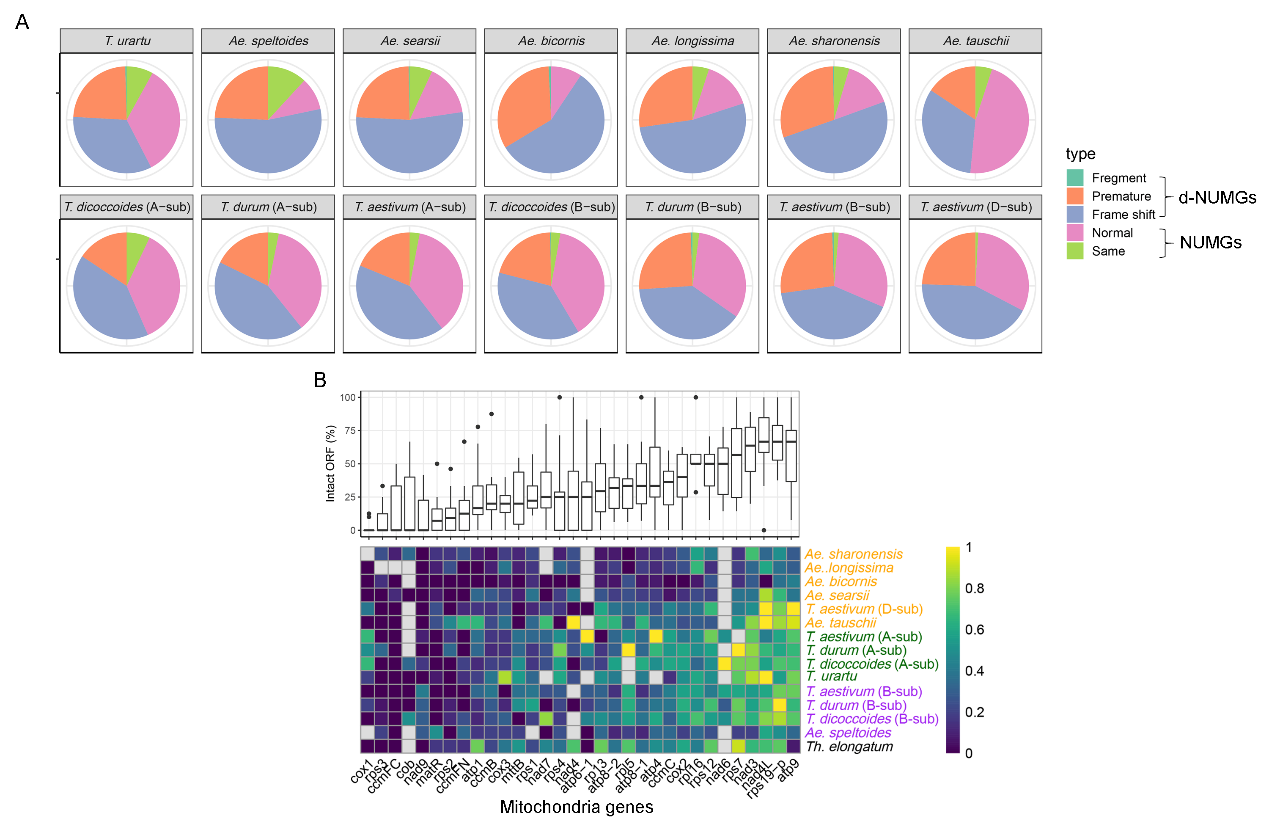

**Figure S5. The genetic fate of genes inside NUMTs. (A)** Different genetic fates of genes within NUMTs. Three groups including lost start and terminal codons (fragment), SNP/InDel induced premature (premature), and frameshift were defined as d-NUMGs (disruption of ORF), whereas those maintained original (same) and larger than 50% amino acid similarity (normal) of mitochondria gene sequences were defined as NUMGs (maintaining of intact ORF). **(B)** The proportion of intact ORFs (NUMGs) for each mitochondria gene among different genomes/subgenomes. The box plot on the top panel shows the median proportion of each mitochondria gene among 13 genomes/subgenomes. The heatmap on the bottom panel gives detail informationfor each mitochondria gene in each genome/subgenome. The corresponding information of NUPTs is shown in **Figure 3**.

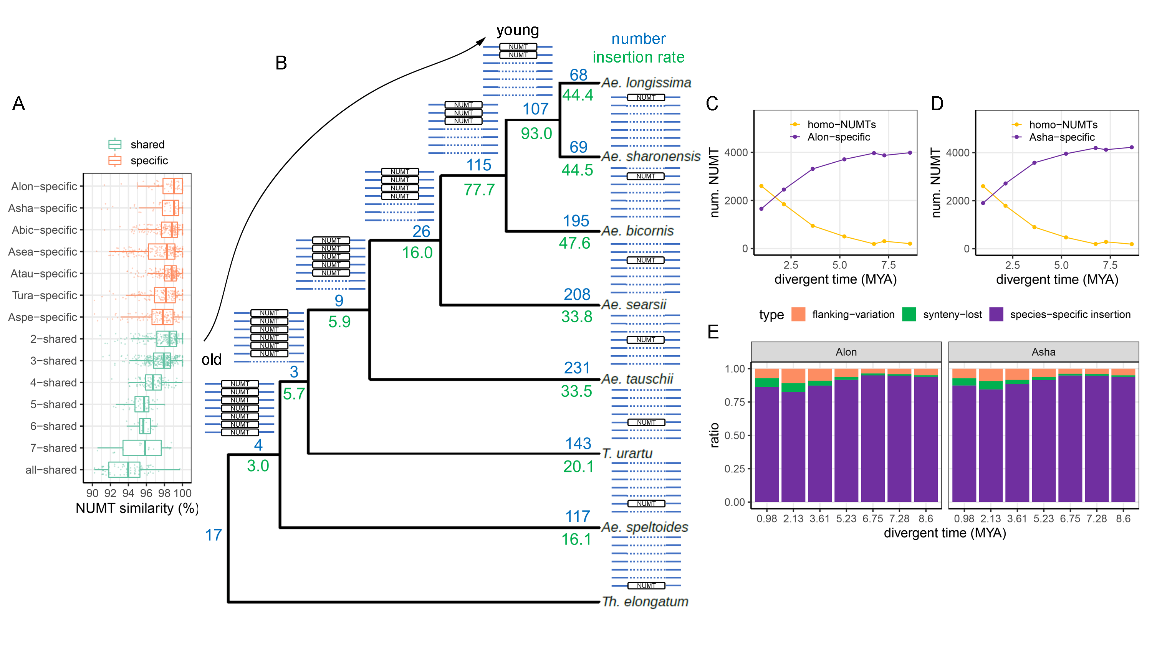

**Figure S6. Evolution of NUMTs during species differentiation among diploids from *Triticum/Aegilops* complex species. (A)** Phylogeny-based statistics of NUMTs. The ideograms of shared and specific homo-NUMT groups were drawn near each node and tip, respectively. Blue and green numbers represent the number and relative insertion ratio (insertion number between two adjacent nodes divided by the evolution time between the corresponding two adjacent nodes) of NUMTs in each node/tip. **(B)** Statistics of NUMT similarity (sequence similarity between NUMT and corresponding DNA fragment in mitochondria genome sequence) for shared and species-specific homo- NUMT groups. **(C) – (D)** Change patterns of homo-NUMT pairs and species-specific NUPTs over divergent time taken *Ae. longissima* **(C)** and *Ae. sharonensis* **(D)** as base (anchor) species, respectively. For each point, the x-axis means the divergent time between the base species and one of the rest species. **(E)** The proportion of different types of non-homo-NUMTs in base (anchor) species when compared to different non-base species. “Flanking variation” means a given NUMT has synteny a counterpart in the non-base species but the flanking regions are not aligned with each other. “Synteny lost” means a given NUMT has a counterpart with flanking regions matched but lost synteny relationship. The corresponding information of NUPTs is shown in **Figure 6**.

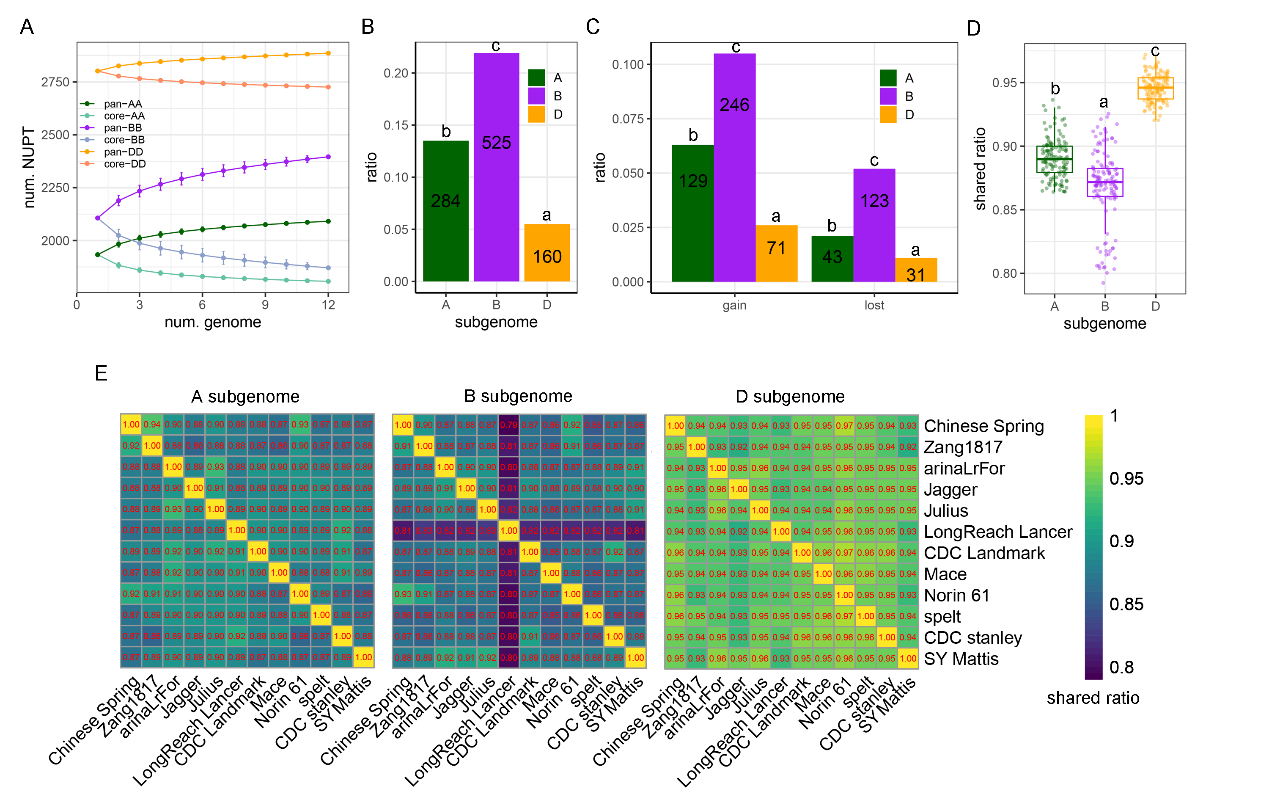

**Figure S7. Subgenomic asymmetry of NUMTs polymorphism among bread wheat accessions. (A)** The profiling of pan-NUMTs and core-NUMTs among 12 hexaploid wheat genomes using pan-genome based analysis according to 2,091 (A-subgenome), 2,396 (B-subgenome), and 2,886 (D-subgenome) homo-NUMT groups. **(B)** Comparisons of NUMT polymorphism ratio ((N_pan-NUMTs_ – N_core-NUMTs_)/ N_pan-NUMTs_) among A-, B- and D-subgenomes. Alphabets mean the results of multiple comparisons based on χ^2^ test and post hoc test. The numbers mean the difference between the numbers of pan-NUMTs and core-NUMTs. **(C)** The proportion of gain and lost homo-NUMT groups among three subgenomes in hexaploidy wheat. Alphabets mean the results of multiple comparisons based on χ^2^ test and post hoc test. The number of gain and lost groups were also shown. **(D)** pairwise comparisons of homo-NUMT pairs among 12 genomes for three subgenomes. The number in each cell represents the proportion of homo-NUMTs to the total number of NUMTs for each comparison. **(E)** Summary of homo-NUMT ratios among three subgenomes based on 12 hexaploid genome datasets. Alphabets mean the results of multiple comparisons based on Kruskal–Wallis rank sum test and Tukey–Kramer test (*p* value < 2.2e-16). The corresponding information of NUPTs is shown in **Figure 8**.
